# WEE1 inhibitors trigger GCN2-mediated activation of the integrated stress response

**DOI:** 10.1101/2025.03.12.642754

**Authors:** Rinskje B. Tjeerdsma, Timothy F. Ng, Maurits Roorda, Daniëlle Bianchi, Sora Yang, Clara Bonnet, Michael VanInsberghe, Marieke Everts, Femke J. Bakker, H. Rolf de Boer, Nathalie Moatti, Nicole Hustedt, Jay Yin, Lisa Hoeg, Matthew Leibovitch, Frank Sicheri, Alexander van Oudenaarden, Steven de Jong, Jeroen van den Berg, Marvin E. Tanenbaum, Thijn R. Brummelkamp, Daniel Durocher, Marcel A.T.M. van Vugt

## Abstract

The WEE1 kinase negatively regulates CDK1/2 to control DNA replication and mitotic entry. Genetic factors that determine sensitivity to WEE1 inhibitors (WEE1i) are largely unknown. A genome-wide insertional mutagenesis screen revealed that mutation of *EIF2A*, a translation regulator, sensitized to WEE1i. Mechanistically, WEE1i treatment triggers a translational shut-down, which is lethal in combination with the reduced translation of *EIF2A*^KO^ cells. A genome-wide CRISPR-Cas9 screen revealed that inactivation of integrated stress response (ISR) kinases GCN1/2 rescued WEE1i-mediated cytotoxicity. WEE1i induced GCN2 activation, ATF4 upregulation, and altered ribosome dynamics. Loss of the collided ribosome sensor ZNF598 conversely increased sensitivity to WEE1i. Notably, the ISR was not required for WEE1i to induce DNA damage, premature mitotic entry or sensitization to DNA-damaging chemotherapeutics. ISR activation was independent of CDK1/2 activation. Importantly, WEE1i-mediated ISR activation was independent of WEE1 presence, pointing at off-target effects, which are shared by multiple chemically distinct WEE1i. This response was also observed in peripheral blood mononuclear cells. Importantly, low-dose WEE1 inhibition did not induce ISR activation, while it still synergized with PKMYT1 inhibition. Taken together, WEE1i triggers toxic ISR activation and translational shutdown, which can be prevented by low-dose or combination treatments, while retaining the cell cycle checkpoint-perturbing effects.

## Introduction

Cell cycle progression is driven by the temporally controlled activation of specific cyclin-dependent kinases (CDKs) in complex with their cognate cyclin partners^1^. Although the roles and regulation of cyclin-CDKs in mammalian cells is complex, distinct roles of specific cyclin-CDKs have been defined; entry into the cell cycle is largely driven by CDK4/6 in complex with cyclin D, initiation and progression of S-phase is promoted by CDK2 in complex with cyclin E and cyclin A, whereas cyclin B1-CDK1 drives entry into mitosis^1^.

An important regulatory mechanism controlling the activity of CDK1 and CDK2 involves inhibitory phosphorylation. Both CDK1 and CDK2 are phosphorylated on Tyr15 (Y15) by the WEE1 kinase^2–4^, which limits their activity and safeguards DNA replication and prevents premature entry into mitosis respectively^5^. In addition, CDK1 is phosphorylated on Thr14 (T14) by the PKMYT1 kinase (also known as Myt1) to prevent unscheduled CDK1 activation^6,7^. To activate CDK1/2, the inhibitory phosphorylation at Y15 and T14 is removed by CDC25 phosphatases^8^. In situations of DNA damage, the DNA damage-regulated kinases ATR and ATM trigger activation of the downstream checkpoint kinases CHK1 and CHK2^9^, which subsequently phosphorylate and inhibit CDC25 phosphatases^8,10^. In parallel, CHK1 activates the WEE1 kinase^11^. Consequently, DNA lesions and incomplete DNA replication lead to maintenance of CDK1/2 inhibitory phosphorylation, which arrests cell cycle progression and allows cells to repair DNA or complete replication.

Inactivation of cell cycle control is cytotoxic in situations of cell-intrinsic or therapy-induced DNA damage^12,13^. This concept is therapeutically exploited by development of a range of cell cycle checkpoint inhibitors, including inhibitors of WEE1^14^. WEE1 inhibitors (WEE1i) were demonstrated to be preferentially cytotoxic in *TP53* mutant cells^14,15^, as well as in cells with elevated levels of replication stress, due to oncogene overexpression^16,17^ or nucleotide deficiency^18^. Moreover, WEE1i treatment has been demonstrated to sensitize tumor cells to a range of genotoxic chemotherapeutic agents^19–22^.

Initial clinical studies focused on combination treatment of the WEE1i adavosertib (AZD1775) in combination with platinum-based chemotherapeutics in patients with high-grade serous ovarian cancer. This combination treatment showed promising results, although dose-limiting toxicities occurred^23–26^. Although all included patients harbored *TP53* mutation tumors, considerable variation in response to WEE1i treatment was observed. Clearly, other tumor-associated features beyond *TP53* mutations influence WEE1i response. In this study we explored genetic determinants of WEE1i treatment response.

## Results

### A haploid genetic screen identifies genes determining sensitivity to the WEE1 inhibitor AZD1775

To identify gene mutations that sensitize cells to inhibition of WEE1, an insertional mutagenesis screen was performed in near-haploid HAP1 cells. To this end, HAP1 cells were mutagenized using a gene trap virus encoding a strong splice acceptor (Suppl. Fig. 1A)^27^, and subsequently treated with the WEE1i AZD1775 (Fig. 1A). Several genes were identified that conferred sensitivity to AZD1775 treatment upon inactivation (Fig. 1B), including multiple genes involved in the one-carbon metabolic pathway, which uses folate to catabolize serine into purines and pyrimidines^28^. Specifically, *MTHFD1, MTHFD1L, MTHFD2,* and *SHMT2* of this pathway were identified (Fig. 1B). Of note, pharmacological targeting of MTHFD2 was previously shown to induce replication stress and increase sensitivity to cell cycle checkpoint inhibitors, including WEE1i^29^. Furthermore, we identified ubiquitin ligase subunits, including the F-box factor *FBXW7* that regulates cyclin E1/CDK2 activity, which was previously described to determine WEE1i sensitivity^16,17,30^. Additionally, inactivation of the PP2A phosphatase activator *PTPA* and *PPP2R5E* (encoding the PP2A subunit B56ε) conferred sensitivity to AZD1775, in line with a previously described role for PP2A in regulating WEE1/CDC25C pathway activation^31^. Identification of these genes underscored the validity of our screening approach.

**Figure 1.**
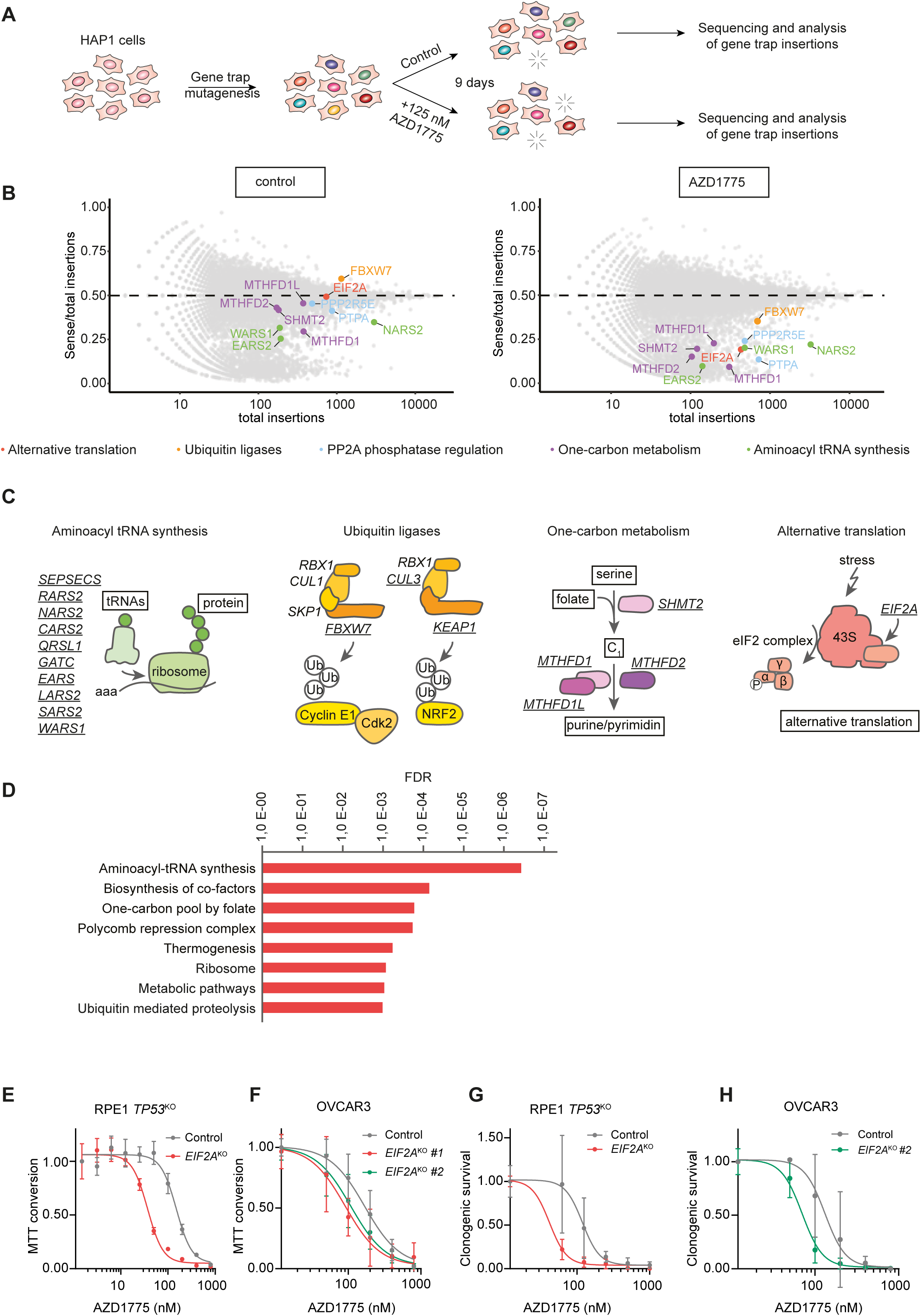
Loss of *EIF2A* sensitizes cells to WEE1 inhibition. **(A)** Experimental setup of the viral insertional mutagenesis screen in. **(B)** Sense/total insertion ratios from mutagenesis screens performed in HAP1 cells treated with AZD1775 or DMSO. Genes with significantly lower sense insertion in AZD1775 treated cells are indicated. **(C)** Schematic representation of high-ranking gene mutations causing sensitization to WEE1i. **(D)** Pathway enrichment analysis, depicting the most highly enriched pathways involved in sensitization to WEE1i. **(E, F)** Dose-response curve of AZD1775 in control RPE1 *TP53*^KO^ and RPE1 *TP53*^KO^ *EIF2A*^KO^ cells (panel E) and OVCAR3, OVCAR3 *EIF2A*^KO^ #1 and OVCAR3 *EIF2A*^KO^ #2 cells (panel F). Cells were treated for 5 days and cellular viability was measured by MTT conversion. Data represent mean ± standard deviation (SD) (n=3). **(G, H)** Dose-response clonogenic survival analysis of control RPE1 *TP53^KO^* and RPE1 *TP53^KO^ EIF2A*^KO^ cells (panel G) and OVCAR3 and OVCAR3 *EIF2A*^KO^ #2 cells (panel H) treated with AZD1775 for 10 days (Representative images are depicted in Suppl. Fig. 1C and 1D, respectively). Data represent mean ± SD (n=2 for RPE1 *TP53*^KO^; n=1, 3 technical replicates for OVCAR3).

Remarkably, we identified multiple genes involved in the regulation of mRNA translation, including many aminoacyl-tRNA synthetases, such as *NARS2*, *RARS2*, *CARS2*, *LARS2*, *SARS2,* and *WARS1* (Fig. 1B-D). Inactivation of these genes mildly affected the viability of control cells but further reduced viability of WEE1-inhibited cells. In contrast, mutation of *EIF2A* did not affect HAP1 cell viability but caused a significant loss of viability upon WEE1 inhibition. The protein eIF2A has been implicated in mRNA translation, specifically in translation of alternative initiation sites^32^. In addition, eIF2A was demonstrated to directly interact with eIF5B^33^ and bind to 40S ribosome subunits^34^, although the exact role of eIF2A herein remains unclear ^35–37^. To confirm its sensitization to WEE1i, *EIF2A* was inactivated using CRISPR/Cas9 in HAP1, RPE1 *TP53*^KO^, and OVCAR3 cells (Suppl. Fig. 1B-F). In a panel of HAP1-*EIF2A* knockout (KO) clones, decreased clonogenic survival was observed when cells were treated with WEE1i (Suppl. Fig. 1B). Likewise, inactivation of *eIF2A* in RPE1 *TP53*^KO^ and OVCAR3 cells conferred WEE1i sensitivity in short-term viability assays (Fig. 1E, F) and clonogenic survival assays (Fig. 1G, H; Suppl. Fig. 1D, F). Combined, these results indicate that loss of eIF2A leads to cellular sensitivity to the WEE1i AZD1775.

### eIF2A loss does not perturb DNA replication

We hypothesized that loss of *EIF2A* may sensitize cells to WEE1 inhibition by increasing DNA replication stress and DNA damage. However, analysis of γH2AX levels did not reveal an increased level of DNA lesions in *EIF2A*^KO^ cells (Fig. 2A, B). In response to WEE1 inhibition, an increase in DNA damage was observed in control and *EIF2A*^KO^ cells (Fig. 2A, B), which included pan-nuclear γH2AX staining reflecting replication stress (Fig. 2C). Although the increase in mean γH2AX intensity in *EIF2A*^KO^ cells was similar to that of control RPE1 *TP53*^KO^ cells, we observed a minor increase in pan-nuclear γH2AX in *EIF2A*^KO^ cells upon WEE1i treatment (Fig. 2C).

**Figure 2.**
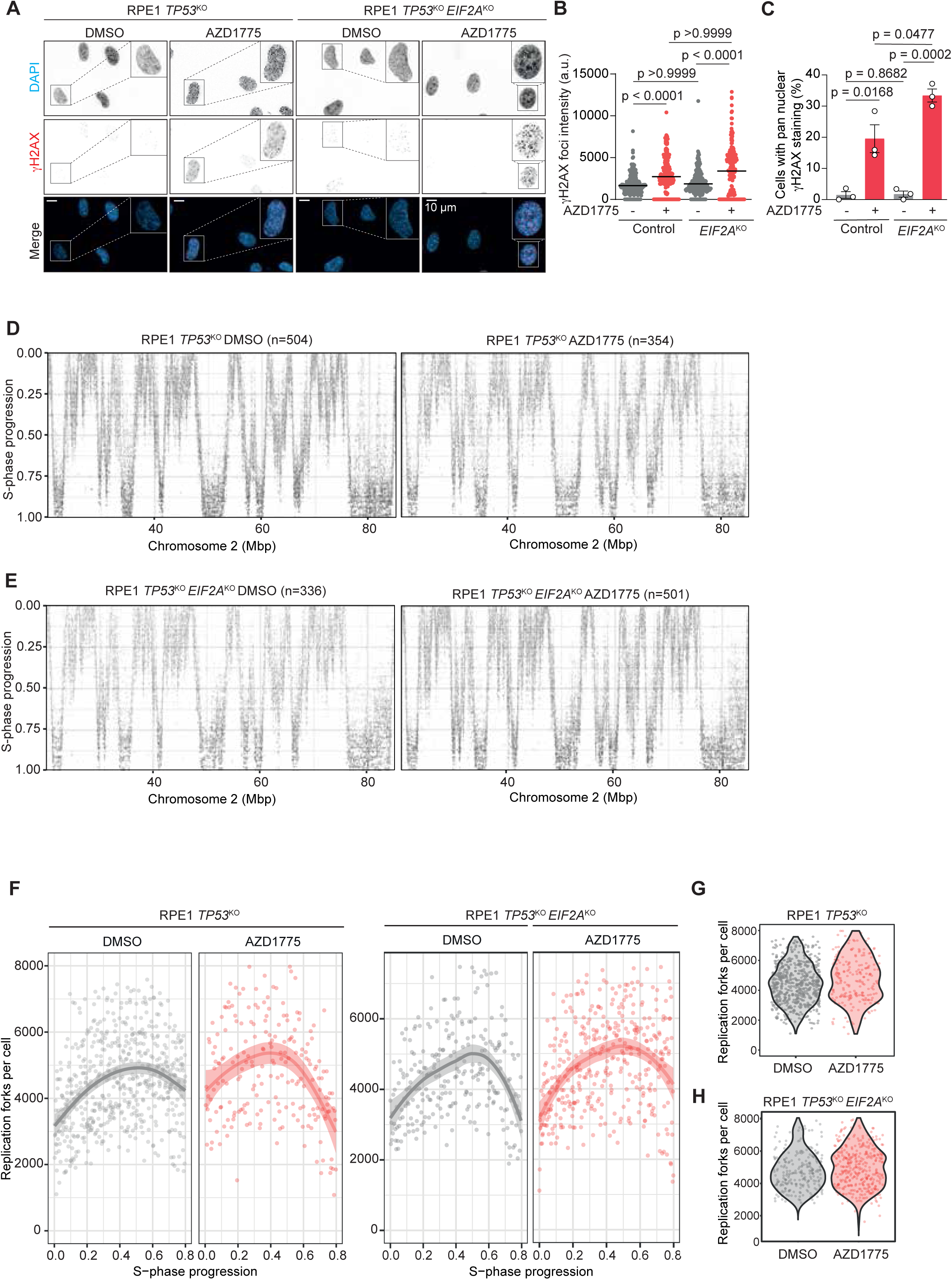
Loss of *EIF2A* does not significantly perturb DNA replication upon WEE1 inhibition. **(A)** Representative immunofluorescence microscopy images of γH2AX staining in control RPE1 *TP53*^KO^ and *EIF2A*^KO^ cells treated with AZD1775 (500 nM) for 24 h. Scalebars indicate 10 µm. **(B)** Mean foci intensity analysis of γH2AX immunofluorescence staining, as shown in panel A. Medians of n=3 experiments, with at least 50 cells counted per experiment for each condition. Kruskal-Wallis ANOVA with Dunn’s multiple comparisons test, p≤0.05 was considered significant. **(C)** Pan nuclear γH2AX immunofluorescence staining analysis of images, as shown in panel A. Median of n=3 experiments ± SEM is indicated, with at least 50 cells counted per experiment for each condition. Unpaired t-test, p≤0.05 was considered significant. **(D)** scEdU-seq maximum normalized log counts are plotted for untreated (left) or AZD1775-treated (125 nM, right) control RPE1 *TP53*^KO^ cells, ranked according to S-phase progression (y-axis) and binned per 400 kb (x-axis). A 40 megabase region of chromosome 2 is shown. **(E)** scEdU-seq maximum normalized log counts were plotted for untreated (left) or AZD1775-treated (125 nM, right) RPE1 *TP53*^KO^ *EIF2A*^KO^cells, ranked according to S-phase progression (y-axis) and binned per 400 kb (x-axis). A 40 megabase region of chromosome 2 is shown. **(F)** Number of DNA replication forks per cell relative to S-phase progression in untreated or AZD1775-treated (125 nM) control RPE1 *TP53*^KO^ (left) or *EIF2A*^KO^ (right) cells, measured by scEdU-seq. **(G)** Number of DNA replication forks per cell in untreated or AZD1775-treated (125 nM) control RPE1 *TP53*^KO^ cells measured by scEdU-seq. **(H)** Number of DNA replication forks per cell in untreated or AZD1775-treated (125 nM) RPE1 *TP53*^KO^ *EIF2A*^KO^ cells measured by scEdU-seq.

Analysis of replication dynamics using single-cell EdU sequencing (scEdUseq) did not reveal major differences between control RPE1 *TP53*^KO^ and *EIF2A*^KO^ cells (Fig. 2D-E left panels, Fig. 2H, Suppl. Fig. 2E). Similarly, AZD1775 treatment did alter replication profiles between control and *EIF2A*^KO^ cells (Fig. 2D-E right panels, Suppl. Fig. 2A). Specifically, the overall number of DNA replication forks per cell were similar between control RPE1 *TP53*^KO^ and *EIF2A*^KO^ cells, as was the distribution of active replication forks over the various stages of S-phase (Fig. 2F-H), leading to a strong positive correlation in replication timing at all stages of S phase between control and *EIF2A*^KO^ cells (Suppl. Fig. 2B). DNA fiber analysis confirmed these findings, showing only marginally decreased DNA track lengths in RPE1 *TP53*^KO^ *EIF2A*^KO^ and OVCAR3 *EIF2A*^KO^ cells when compared to control cells (Suppl. Fig. 2C). These data indicate that *EIF2A*^KO^ cells do not have a significant increase in replication stress levels.

Treatment with AZD1775 resulted in an expected and dose-dependent perturbation of cell cycle dynamics, causing a higher proportion of G2-phase and mitotic cells and with fewer S-phase cells (Suppl. Fig. 2D). Low-dose (125 nM) AZD1775 did not dramatically alter replication dynamics (Fig. 2D, F, H), but we could detect a mild increase in replication forks at S-phase onset, followed by a decrease during S-phase progression (Fig. 2F). At a high-dose of AZD1775 treatment (1 µM), DNA replication was dramatically perturbed (Suppl. Fig. 2E-G), which coincided with a strong decrease in active replication forks per cell (Suppl. Fig. 2F, G). Importantly, no striking differences in replication dynamics were observed in *EIF2A*^KO^ cells after low dose AZD1775 treatment (Fig. 2E, H), at which dose *EIF2A*^KO^ cells already showed increased AZD1775 sensitivity (Fig. 1E, G). Taken together, these data suggest that neither alteration in the DNA replication program not replication stress are driving the cytotoxic effects of WEE1 inhibition in *EIF2A*^KO^ cells.

### WEE1 inhibition induces the integrated stress response through GCN2 activation

Comparison of baseline mRNA expression levels between control RPE1 *TP53*^KO^ and *EIF2A*^KO^ cells revealed significant downregulation of gene sets related to mRNA translation, including ‘RNA metabolism’, ‘rRNA processing’, and ‘Translation’ (Fig. 3A, Suppl. Fig. 3A, Suppl. Table 1), in line with a proposed regulatory role for eIF2A in mRNA translation^38,39^. Proteomics analysis using stable isotope labeling by amino acids in cell culture (SILAC) confirmed loss of eIF2A protein in *EIF2A*^KO^ cells, along with alteration of translation regulators, including higher protein levels of the translation elongation factor EEF1E1 and the regulator of ribosomal translation LARP1^40–42^ (Fig. 3B, C). Using puromycin or L-azidohomoalanine (AHA) incorporation assays to measure ongoing mRNA translation, we observed that two chemically distinct WEE1i, AZD1775 and Debio 0123, decreased ongoing mRNA translation in RPE1 *TP53*^KO^ cells (Fig. 3D, E). Importantly, mRNA translation attenuation upon WEE1i-treatment was even more pronounced in *EIF2A*^KO^ cells (Fig. 3D). Downregulation of translation upon WEE1i treatment was confirmed in HCC38 triple-negative breast cancer cells (Suppl. Fig. 3B), which was rescued by inhibition of the integrated stress response (ISR) using ISRIB^43^ (Fig. 3E). To provide insights into how WEE1 inhibition affects mRNA translation, we performed a CRISPR-based screen to identify gene mutations that confer resistance to AZD1775 in RPE1 hTERT *TP53*^KO^ cells. (Fig. 3F). The strongest hits included the eIF2α kinase GCN2 (*EIF2AK4*) and its coactivator GCN1 (*GCN1L1*), which function in the integrated stress response, as well as negative mTORC1 regulators TSC1 and TSC2 (Fig. 3G)^44^. To corroborate these results, RPE1 *GCN2*^KO^ and *GCN1*^KO^ cells were generated and WEE1i sensitivity was analyzed. Indeed, both *GCN2*^KO^ and *GCN1*^KO^ cells were less sensitive to AZD1775 treatment compared to RPE1 control cells in short-term proliferation assays (Fig. 3H), clonogenic survival assays (Suppl. Fig. 3C, D) and cell competition assays (Suppl. Fig. 3E, F).

**Figure 3.**
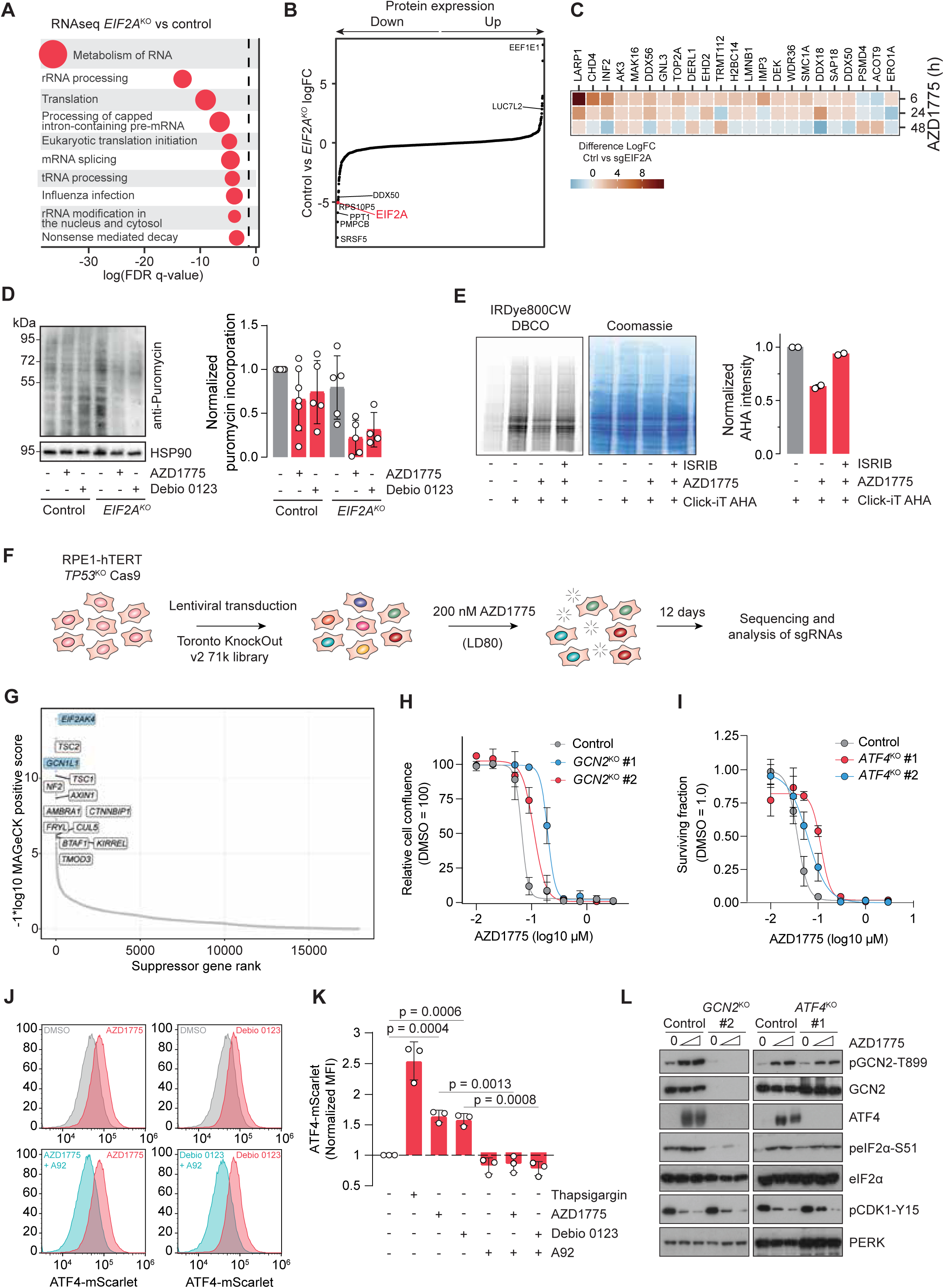
WEE1 inhibition induces a translational attenuation mediated by GCN2 and the integrated stress response. **(A)** Gene set enrichment analysis of RNA-seq data of RPE1 *TP53*^KO^ control and *EIF2A*^KO^ cells (more detailed analysis in Suppl. Fig. 3A). Circle size represents gene set size. **(B, C)** SILAC-based proteomics analysis of RPE1 *TP53*^KO^ control and *EIF2A*^KO^ cells, showing differential protein expression at baseline (panel B) and upon AZD1775 treatment for 6, 24 and 48 h (panel C). **(D)** RPE1 *TP53*^KO^ control and *EIF2A^KO^* cells were treated with AZD1775 or Debio 0123 (1 µM) for 24 h, followed by a 10’ puromycin pulse. Puromycin incorporation was visualized by immunoblotting (left panel). Quantification of immunoblots is indicated (right panel). Data represent mean ± SD; RPE1 *TP53*^KO^ *EIF2A*^KO^ treated with Debio 0123 (n=4); control RPE1 *TP53*^KO^ treated with DMSO or AZD1775 (n=7); Other conditions (n=5). **(E)** RPE1 *TP53*^KO^ *PAC*^KO^ cells were treated with AZD1775 (250 nM) and ISRIB for 24 h and subsequently labeled with L-azidohomoalanine (AHA). Immunoblot of AHA-labeled proteins and quantification is shown. Data represent mean ± SD (n=2). **(F)** Schematic overview of the CRISPR/Cas9 genome-wide AZD1775 resistance screen. **(G)** MAGeCK scores for individual genes in the RPE1 *TP53*^KO^ AZD1775 positive selection screen (day 12 vs day 0). **(H)** Cell confluency analysis of parental RPE1 *TP53^KO^ PAC*^KO^ or *GCN2^KO^* cells. Mean ± SD, n=3 for control and *GCN2*^KO^ #1, n=2 for *GCN2*^KO^ #2. **(I)** Quantification of clonogenic survival assays of parental RPE1 *TP53*^KO^ *PAC*^KO^ and *ATF4^KO^* #1 and #2 cells (corresponding images provided in Suppl. Fig. 3H). Mean ± SD, n=3 for all conditions. **(J, K)** Representative histograms (panel J) and quantification (panel K) of ATF4-mScarlet flow cytometry measurements in RPE1 *TP53*^KO^ ATF4-mScarlet reporter cells treated with thapsigargin (1 µM), AZD1775 (1 µM), Debio 0123 (1 µM) and/or A92 (1 µM) for 24 h. Data represent mean ± SD (n=3). **(L)** Parental RPE1 *TP53*^KO^ *PAC*^KO^, *ATF4*^KO^ #1 and *GCN2*^KO^ #2 cells were treated with AZD1775 (0, 250 or 500 nM) for 24 h, and indicated proteins were immunoblotted. Statistical analysis of panels D, K: unpaired t-test with p ≤ 0.05 considered significant.

Subsequently, we analyzed whether signaling downstream of GCN2 also determined AZD1775-induced cytotoxicity. Time course analysis of AZD1775-induced activation of GCN2 and the downstream transcription factor ATF4^45^ showed GCN2 phosphorylation and ATF4 upregulation already at 30 minutes after AZD1775 treatment (Suppl. Fig. 3G). Inactivation of ATF4 led to reduced sensitivity of RPE1 *TP53*^KO^ cells to AZD1775 (Fig. 3I, Suppl. Fig. 3H). These findings were confirmed in RPE1 *TP53*^KO^ cells harboring a fluorescent ATF4 reporter, consisting of two upstream open reading frames (uORFs) and the first 84 base pairs of the *ATF4* gene (ORF3) fused to an *mScarlet* cassette^46^. Under unstressed conditions, only the two uORFs are translated, whereas upon ISR activation, delayed translation reinitiation leads to ribosome trailing past the initiation site of uORF2 and translation of the ATF4-mScarlet fusion protein^46^. Thapsigargin, a known activator of the ISR via the PERK kinase, confirmed the ability of these reporter cell lines to read out ATF4 upregulation (Suppl. Fig. 3I, J). Treatment with AZD1775 or the chemically distinct WEE1i Debio 0123 led to ATF4-mScarlet upregulation, which was fully rescued by pharmacological GCN2 inhibition (Fig. 3J, K, Suppl. Fig. 3I, J). Supporting this finding, pharmacological inhibition of GCN2 also rescued AZD1775-mediated reduction of mRNA translation in the breast cancer cell line HCC38 (Suppl. Fig. 3B). Furthermore, inactivation of *GCN2* prevented AZD1775-induced ATF4 upregulation and eIF2α phosphorylation, but did not affect CDK1-Y15 phosphorylation, whereas ATF4 loss only impacted ATF4 levels as it functions downstream of eIF2α-S51 phosphorylation and the suppression of cap-dependent translation (Fig. 3L). Taken together, these data show that WEE1i trigger activation of the ISR and decrease global mRNA translation in a GCN2-dependent manner.

### WEE1i-mediated induction of the ISR is cell cycle-independent

Since loss of ATF4 or GCN2 did not affect CDK1-Y15 phosphorylation (Fig. 3L), we analyzed whether GCN2 activation by WEE1 inhibition is dependent on cell cycle status. Quantitative image-based cytometry (QIBC) showed that ATF4 expression was upregulated upon AZD1775 treatment in RPE1 *TP53*^KO^ cells throughout the cell cycle, which was prevented by ISRIB co-treatment (Fig. 4A). To further investigate whether ATF4 upregulation by AZD1775 requires ongoing cell cycle progression, RPE1 *TP53*^KO^ were synchronized in mitosis using the microtubule-poison nocodazole, and released into a round of cell division. Alternatively, cells were arrested in G1 phase after release from mitosis, using the CDK4/6 inhibitor palbociclib (Fig. 4B). Subsequently, cells were treated with AZD1775 and/or the GCN2 inhibitor A92. A palbociclib-mediated arrest in G1 was confirmed by low abundance of cyclin B (Suppl. Fig. 4A). AZD1775 treatment resulted in upregulation of ATF4, both in proliferating cells as well as G1-arrested cells (Fig. 4C, Suppl. Fig. 4A, B). Conversely, GCN2 inhibition prevented ATF4 upregulation in both cycling and non-cycling cells, indicating that the GCN2-mediated ATF4 upregulation in response to WEE1 inhibition is independent of cell cycle status.

**Figure 4.**
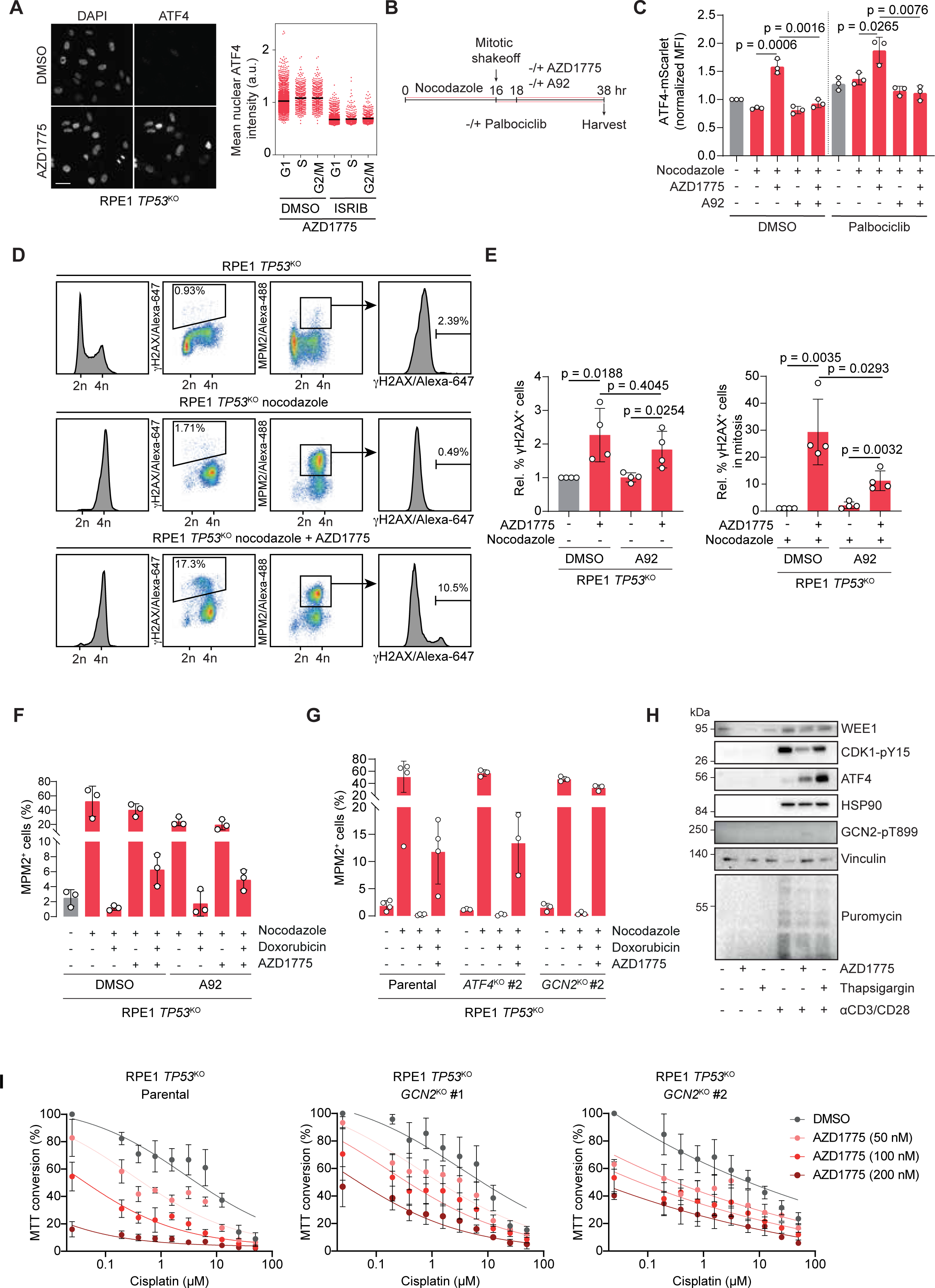
Integrated stress response activation by WEE1i is cell cycle independent. **(A)** RPE1 *TP53*^KO^ *PAC*^KO^ cells were treated with AZD1775 (250 nM) for 6 h in the presence or absence of ISRIB (1 μM), pulse labeled with EdU, and processed for quantitative image-based cytometry. Cell cycle stage was defined by DNA content and EdU positivity. Median ATF4 intensity from n=2 experiments is shown. **(B, C)** Experimental set-up (panel B). RPE1 *TP53*^KO^ ATF4-mScarlet-NLS reporter cells were treated overnight with nocodazole. Mitotic cells were isolated and replated in the presence or absence of palbociclib. After 2 h AZD1775 (1 µM) and/or A92 (1 µM) was added for 20 h (Panel B). mScarlet levels in RPE1 *TP53*^KO^ ATF4-mScarlet reporter cells were measured by flow cytometry. Data represent mean ± SD (n=3)(panel C). **(D, E)** Flow cytometry gating strategy (panel D) and analysis of γH2AX in RPE1 *TP53^KO^* cells after treatment with AZD1775 (1 µM) and/or A92 (1 µM) in the absence (panel E, left) or presence (panel E, right) of nocodazole. Data represent mean ± SD (n=4). **(F)** Flow cytometry analysis of MPM2-positivity in RPE1 *TP53^KO^* cells after treatment with nocodazole, doxorubicin, AZD1775 (1 µM) and/or A92 (1 µM). Data represent mean ± SD (n=3). **(G)** Flow cytometry analysis of MPM2-positivity in control RPE1 *TP53*^KO^ *PAC*^KO^ and *ATF4*^KO^ #2 or *GCN2*^KO^ #2 cells after treatment with nocodazole, doxorubicin and AZD1775 (1 µM). Data represent mean ± SD (n=3). **(H)** Immunoblot of resting PBMCs or PBMCs stimulated with anti-CD3/CD28 beads. PBMCs were treated with DMSO, AZD1775 (500 nM) or thapsigargin (500 nM) for 24 h. **(I)** RPE1 *TP53*^KO^ *PAC*^KO^ and *ATF4*^KO^ #2 or *GCN2*^KO^ #2 cells were treated with AZD1775 and/or cisplatin for 5 days. Cell survival was analyzed by MTT conversion. Data represent mean ± SD (n=4). Statistical analysis was performed using unpaired t-tests, with p ≤ 0.05 considered significant.

WEE1i are well described to cause DNA damage and premature entry into mitosis^12,47^. Since loss of GCN2 reduced WEE1i-induced cytotoxicity, we analyzed whether GCN2 was required for WEE1i-induced DNA damage and premature mitotic entry. As expected, WEE1 inhibition significantly increased the percentage of γH2AX-positive cells, which is caused by unscheduled cleavage of replication forks^48^, a phenotype that was not rescued by GCN2i (Fig. 4D; 4E, left panel). WEE1 inhibition induced entry of γH2AX-positive cells in mitosis, which also occurred in GCN2-inhibited cells (Fig. 4D; 4E, right panel). Of note, GCN2i partially suppressed the entry of γH2AX-positive cells into mitosis, which may have been caused by slower proliferation of GCN2i-treated cells, a possibility supported by the observation that the total number of mitotic cells was lower upon chemical GCN2 inhibition (Suppl. Fig. 4C).

WEE1i can also force entry into mitosis in cells treated with DNA-damaging chemotherapeutic agents^20,49^. Indeed, WEE1 inhibition resulted in an override of the G2 checkpoint arrest induced by the DNA topoisomerase II inhibitor doxorubicin, which was not prevented by GCN2i (Fig. 4F, Suppl. Fig. 4D). To confirm these findings, we analyzed the proportion of mitotic cells upon AZD1775 treatment in the RPE1 *TP53*^KO^ *GCN2*^KO^ or *ATF4*^KO^ cell lines. Neither inactivation of *GCN2* nor *ATF4* affected the ability of AZD1775 to override the doxorubicin-induced G2 cell cycle arrest (Fig. 4G, Suppl. Fig. 4E, F). Taken together, these results show that a WEE1i-mediated cell cycle checkpoint override is independent of ISR activation.

To investigate whether WEE1 inhibition also leads to ISR activation in non-transformed cells, peripheral blood mononuclear cells (PBMCs) were treated with AZD1775 or thapsigargin as a positive control. Non-stimulated PBMCs were compared with PBMCs that were activated using anti-CD3/CD28 T-cell activator beads (Fig. 4H). Ongoing mRNA translation levels were below detection level in unstimulated PBMCs, along with very low levels of WEE1 protein expression (Fig. 4H). In contrast, activated PBMCs showed an overall increase in protein abundance and ongoing mRNA translation (Fig. 4H, Suppl. Fig. 4G). WEE1i treatment in activated PBMCs robustly induced ATF4 expression, and suppressed active mRNA translation (Fig. 4H, Suppl. Fig. 4G). In line with the observed ATF4 induction, GCN2-T899 phosphorylation was increased upon WEE1i treatment (Fig. 4H, Suppl. Fig. 4H). These data show that ISR activation by WEE1i is not limited to immortalized or transformed cells and can occur in human primary cells.

AZD1775 has been evaluated in combination with platinum-based chemotherapeutic agents^23–26^. We analyzed whether GCN2 activation is required for the potentiating effects of AZD1775 on cisplatin sensitivity. Loss of GCN2 partially rescued the cytotoxicity induced by AZD1775 as a single agent, but not the additive cytotoxicity of combined WEE1i and cisplatin treatment (Fig. 4I, Suppl. Fig. 4I). These findings are consistent with the observation that GCN2 inactivation did not prevent γH2AX induction and cell cycle checkpoint override following AZD1775 treatment (Fig. 4E-G). Combined, these data show that activation of GCN2 by WEE1i is cell cycle-independent, and that GCN2 loss does not prevent WEE1i-induced DNA damage or premature mitotic entry.

### WEE1 inhibition alters translation kinetics

Since WEE1i activate the ISR via GCN2, independently of cell cycle status or DNA damage induction, we tested if WEE1i directly affected ribosome dynamics. A CRISPR-based negative selection screen predicted synergistic WEE1i interactors, including *PKMYT1*, suggesting redundancy in the regulation of CDK inhibition, and confirming previous findings^50,51^ (Fig. 5A, B). Moreover, loss of the DNA polymerase epsilon subunits *POLE3* and *POLE4,* as well as the ribonucleotide reductase catalytic subunit *RRM1* sensitized cells to WEE1i. Loss of these genes was previously described to sensitize cells to ATR inhibition, emphasizing the shared roles of WEE1 and ATR in guarding faithful S-phase progression^52^. We also identified *WEE1*, pointing to drug-induced genetic insufficiency, possibly involving inactivation of one *WEE1* allele, rendering cells more sensitive to WEE1i treatment^53^. Finally, we identified a subset of genes that function in ribosome quality control (RQC), including *ZNF598*, *PELO*, *HBS1L*, *USP9X*, *RNF25*, and components of the ASC-1 complex (*ASCC2* and *ASCC3*)^54–58^ (Fig. 5B-C). These observations suggest that mechanisms that resolve ribosome collisions are important to determine WEE1i sensitivity, with *GCN2* and *GCN1* loss appearing as drivers of WEE1i resistance in this screen (Fig. 5B). Sensitization to WEE1i upon inactivation of *ZNF598* was confirmed in Incucyte cell proliferation assays (Suppl. Fig. 5A).

**Figure 5.**
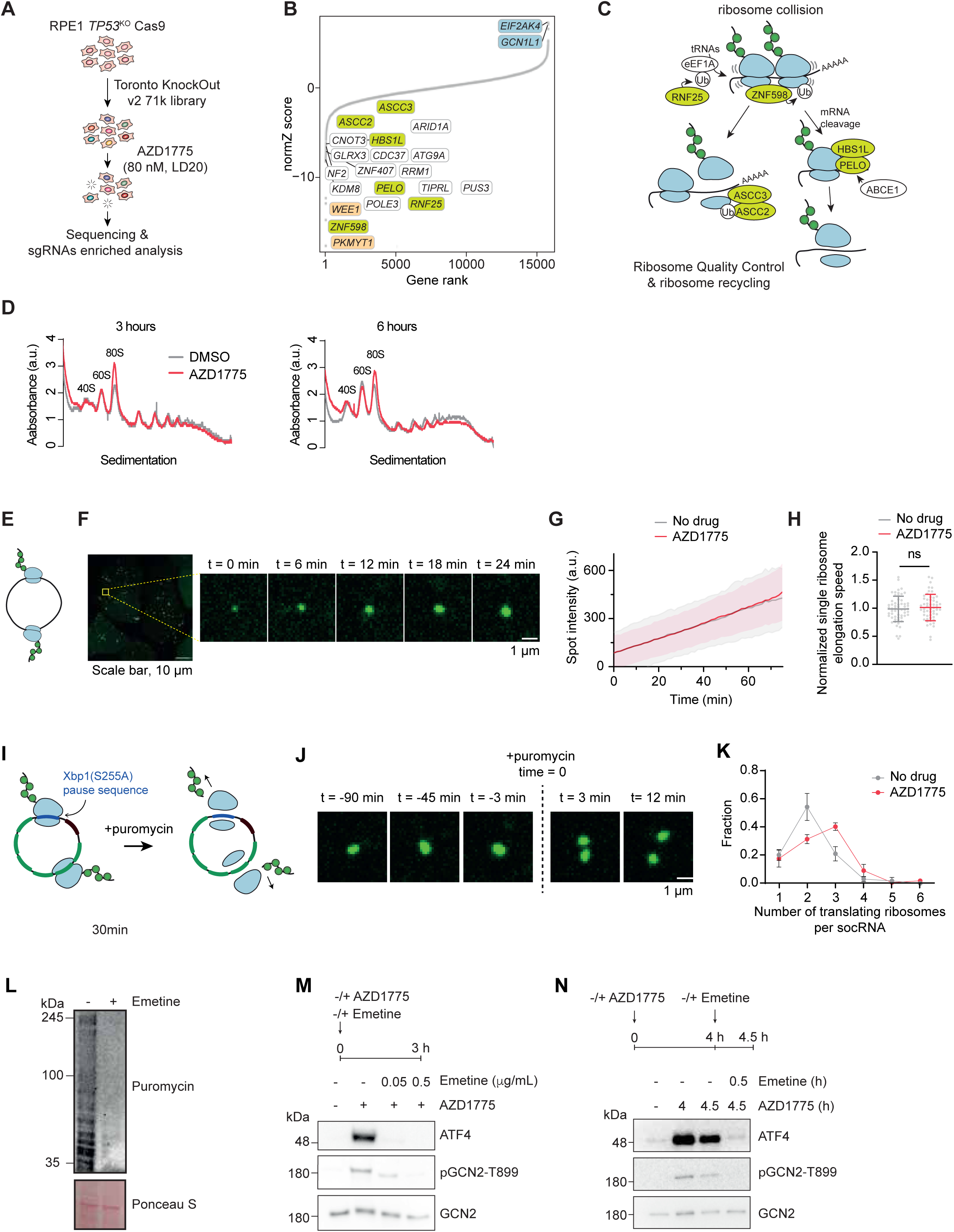
Effects of WEE1i on translation dynamics. **(A)** Schematic overview of the CRISPR screen performed in panel B. **(B)** NormZ scores of individual genes in an RPE1 *TP53*^KO^ *PAC*^KO^ Cas9 negative selection AZD1775 CRISPR screen (day 15 vs. day 0) analyzed by the DrugZ algorithm. Positive normZ scores correspond to suppression and negative normZ scores correspond to synergy. Genes are color-coded by category: orange: cell cycle checkpoint kinases, green: ribosome quality control, blue: ISR kinases *GCN1L1 (*GCN1) and *EIF2AK4* (GCN2). **(C)** Schematic representation of ribosome quality control pathway members with the high-ranking screen hits sensitizing to WEE1i. **(D)** Polysome profiling of RPE1 *TP53*^KO^ cells after treatment with AZD1775 (250 nM) for 3 or 6 h (representative plot of n=2). **(E)** Schematic representation of single ribosome mRNA translation kinetics measured by socRNA-mediated GFP translation. **(F)** Representative image of socRNA-mediated GFP translation. **(G)** GFP foci intensity over time in control or AZD1775-treated conditions. **(H)** Elongation speed of a single ribosomes on a socRNA in control or AZD1775 treated cells (n=2 independent experiments). **(I)** Schematic representation of a socRNA containing an Xbp1 pause sequence. Addition of puromycin triggers the release of ribosomes and nascent peptide chains from a socRNA, allowing the quantification of ribosome copies per socRNA. **(J)** Visual representation of puromycin addition to socRNA. **(K)** Quantification of the number of translating ribosomes per socRNA after 24 h of treatment with AZD1775. Mean ± SD (No drug n=2; AZD1775 n=3). **(L)** Immunoblot of RPE1 *TP53*^KO^ cells after 30 min treatment with emetine (0.5 µg/ml) and puromycin pulse. **(M)** Immunoblot of RPE1 *TP53*^KO^ cells treated with AZD1775 (250 nM) and emetine (0.5 µg/ml) according to the indicated timeline.

These observations led us to investigate whether WEE1i might influence ribosome dynamics. To this end, polysome profiling was performed and revealed elevated 80S ribosome subunit levels on mRNAs upon WEE1i treatment (Fig. 5D). To investigate translation dynamics in more detail, we used the *Stopless-ORF circular* (soc)RNA assay, a recently-developed single-molecule imaging approach to visualize translation dynamics with high precision^59,60^ socRNAs encode SunTag peptide epitopes, which are co-translationally labeled by stably expressed STAb-GFP, resulting in local accumulation of GFP at socRNAs, allowing real-time measurements of translation^59^ (Fig. 5E, F). No difference in ribosome elongation speed was observed between WEE1i-treated cells and untreated cells (Fig. 5G, H). In addition, the number of translating ribosomes per socRNA was not different in cells treated with WEE1i (Suppl. Fig. 5B), indicating that ribosome processivity was also unaffected^60^. To specifically examine ribosome quality control in WEE1i-treated cells, we next analyzed a socRNA harboring the well-established Xbp1(S255A) ribosome pausing sequence. Ribosomes will stall at this sequence, causing ribosome collisions, which will eventually be recycled in a ZNF598-dependent manner^58,61^. In this assay, collision-induced ribosome recycling efficiency can be assessed by measuring the number of ribosomes per Xbp1(S255A) socRNA, which can be determined by inducing ribosome release from the socRNA through puromycin treatment and counting the number of GFP foci (each representing a single ribosome) that split from a single socRNA^60^ (Fig. 5I, J). When compared to control-treated cells, WEE1i-treated cells contained more ribosomes per Xbp1(S255A) socRNA, suggesting that collision-induced ribosome recycling was suppressed in WEE1i-treated cells, with an effect size similar to that reported for ZNF598 depletion (Fig. 5K)^60^. These effects on ribosome recycling were not due to a change in pause duration upon WEE1i treatment (Suppl. Fig. 5C). Defective ribosome recycling after WEE1 inhibition may cause long-lived ribosome collisions in these cells, which results in GCN2 activation, explaining the genetic interaction of WEE1 inhibition with proteins involved in ribosome quality control.

The observed defective ribosome recycling suggested that GCN2 responds to translation perturbations caused by the WEE1i. To test this concept more directly, we asked whether ongoing mRNA translation was necessary for the activation of GCN2 by WEE1i. To do so, we treated RPE1 *TP53*^KO^ cells with the translation inhibitor emetine^62^ at a dose that caused ribosome stalling (Fig. 5L). Notably, we observed that co-treatment of cells with emetine and WEE1i completely blocked GCN2 activation (Fig. 5M). We also observed that emetine addition to cells that undergo ISR caused by WEE1i suppressed GCN2 activation (Fig. 5N), indicating that ongoing translation is required for sustained GCN2 activation by WEE1i treatment.

To further dissect how WEE1i alter ribosome processivity, mRNA-seq and Ribo-seq of AZD1775-treated RPE1 *TP53*^KO^ cells was performed. Differential expression analysis of bulk mRNA-seq data revealed a robust ISR and hallmarks of GCN2 activation, characterized by increased expression of amino acid biosynthesis (e.g. asparagine synthetase (*ASNS*), cystathionine beta-synthase (*CBS*), and phosphoserine aminotransferase 1 (*PSAT1*)) along with aminoacyl-tRNA synthetases (Suppl. Fig. 6A). Robust expression of *DDIT4*, *REDD1* and *SESN2* was observed as well, suggesting that WEE1i mediates mTORC1 inhibition in a GCN2-dependent manner. Moreover, gene set enrichment analysis identified robust upregulation of pathways related to mRNA translation at late time-points of AZD1775 treatment, specifically involving eukaryotic translation initiation and elongation, peptide chain elongation, cap-dependent translation initiation and seleno-amino acid metabolism in response to 48 h AZD1775 treatment (Suppl. Fig. 6B).

Ribosome profiling (Ribo-seq) analysis showed that high-dose WEE1i treatment for 24 h resulted in a global decrease in translation, independently of mRNA abundance (Fig. 6A, B). Pathway enrichment analysis showed that high-dose WEE1i led to significant downregulation of translation-related pathways such as “eukaryotic translation elongation” and “seleno-amino acid metabolism” (Fig. 6C). Upregulated transcription of a panel of ATF4 target genes (i.e. *DDIT3, DDIT4*) (Fig. 6D), as well as amino acid biosynthesis factors such as *ASNS* and *PSAT1* (Suppl. Fig. 6A, B) confirmed that the ISR was activated in these experiments. Although not statistically significant, likely due to limited sample size, the levels of the ATF4 target genes *ATF3* and *PPP1R15A* also appeared to increase (Fig. 6D). In Fig. 6E, ribosome codon occupancy per codon is depicted. Codon occupancy was only mildly shifted after 24 h of WEE1i treatment, possibly due to a shift in the genes expressedas a result of the induction of the alternative translation program by ISR activation, and overall lower coding DNA sequence (CDS) reads (Fig. 6E, Suppl. Fig. 6C, D), and overall translation extension kinetics were similar between conditions (Fig. 6E), in line with our observations of socRNA ribosome kinetics (Fig. 5G). Overall, these data suggest that WEE1i alter translation kinetics and affects ribosome processivity triggering cellular adaptation.

**Figure 6.**
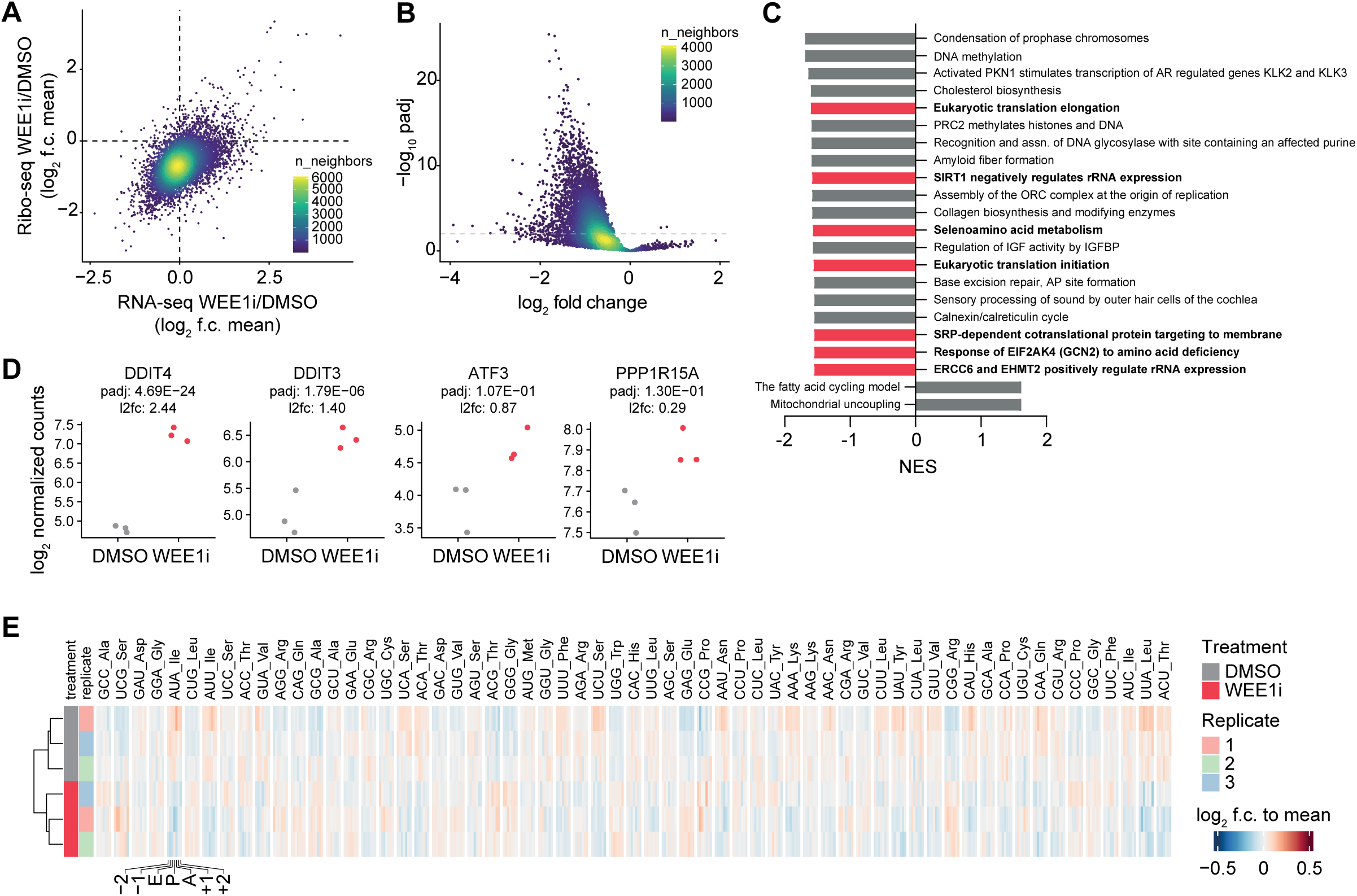
AZD1775 induces wide-spread altered pathway expression but does not affect codon usage. **(A)** Relative gene and protein expression upon treatment with AZD1775 (24 h, 1 µM) measured by Ribo-seq and RNA-seq. **(B)** Differential protein expression upon treatment with AZD1775 (24 h, 1 µM) measured by Ribo-seq. **(C)** Pathway enrichment analysis of Ribo-seq data shown in (M). **(D)** Gene expression of ATF4-target genes *DDIT4, DDIT3, ATF3* and *PPP1R15A* upon treatment with AZD1775 (24 h, 1 µM). **(E)** Codon usage in RPE1 *TP53*^KO^ cells upon treatment with DMSO or AZD1775 (24 h, 1 µM).

### The WEE1i AZD1775 activates GCN2 as an off-target activity

To analyze whether inhibitors of other cell cycle checkpoint kinases also activate the ISR, we measured ATF4-mScarlet upregulation upon treatment of RPE1 *TP53*^KO^ ATF4 mScarlet reporter cells with inhibitors of PKMYT1 (RP-6306), ATR (VE-822), and CHK1 (AZD7762). From this panel, only AZD1775 treatment resulted in clear ATF4-mScarlet upregulation (Fig. 7A).

**Figure 7.**
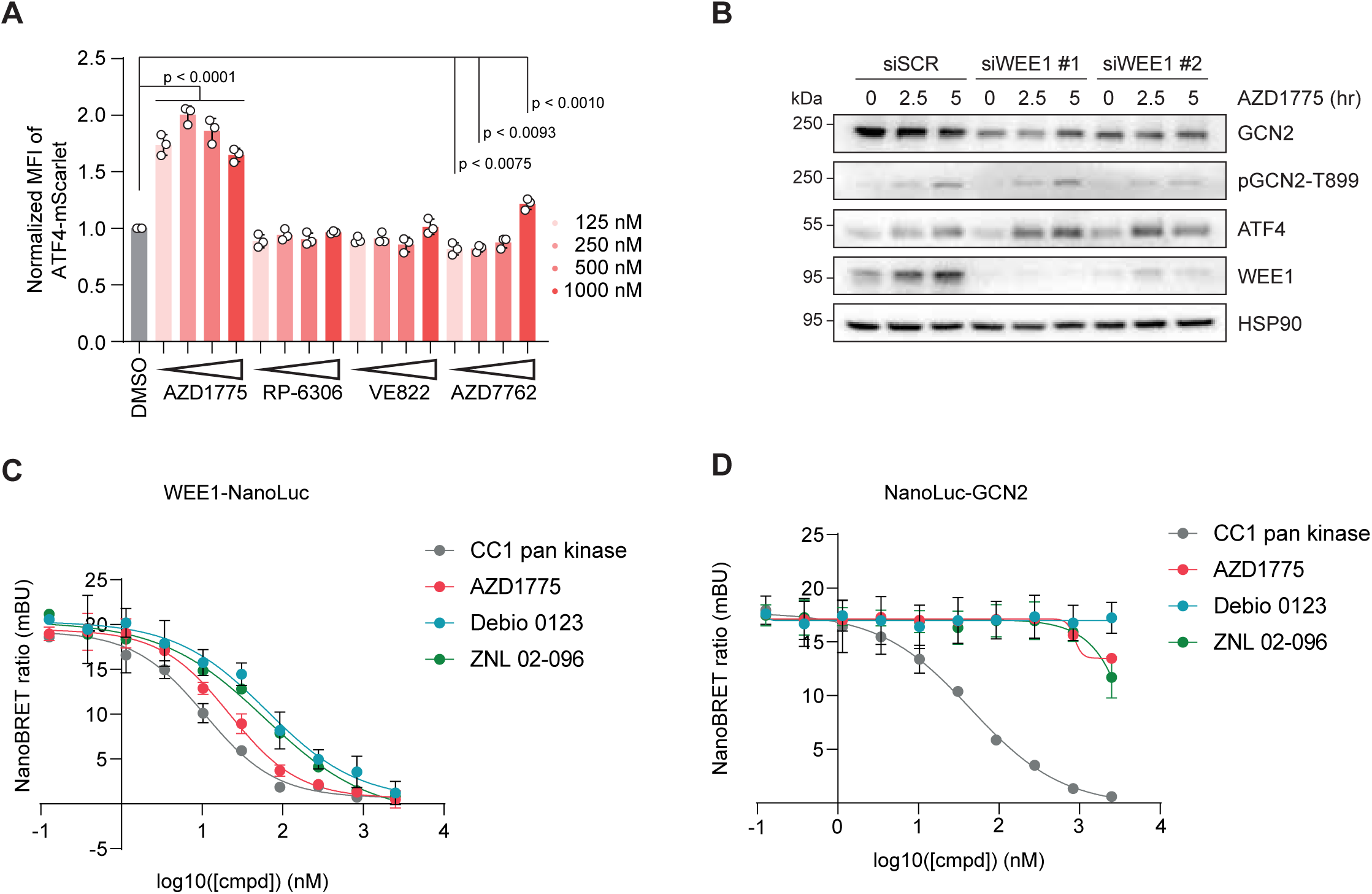
AZD1775 and Debio 0123-mediated GCN2 activation is independent of WEE1. **(A)** Relative ATF4-mScarlet MFI in RPE1 ATF4-mScarlet reporter cells after treatment with 125, 250, 500, or 1000 nM of AZD1775, RP-6306, VE-822 or AZD7762. Data represent mean ± SD (DMSO: n=4; other conditions: n=3). Statistical analysis was performed using a one-way ANOVA multiple comparisons with p≤0.05 considered significant. **(B)** Immunoblot of RPE1 *TP53*^KO^ cells treated with siRNA targeting *WEE1* for 72 h and AZD1775 (250 nM) for 2.5 or 5 h. **(C, D)** NanoBRET kinase target engagement assay performed with HEK293T cells transfected with WEE1-NanoLuc (panel C) and NanoLuc-GCN2 (panel D) and incubated with K-10 tracer (0.5 μM) and indicated doses of CC1, AZD1775, Debio 0123 and ZNL-02-096 for 2 h prior to substrate addition and BRET signal detection.

Small molecule inhibitors of eIF2α kinases, PERK and PKR, as well as tyrosine kinase inhibitors including neratinib and dovitinib have been shown to directly activate GCN2^63,64^. The proposed model for GCN2 activation by these drugs involves binding to the ATP binding pocket of one of the two GCN2 monomers, thereby enhancing the affinity of the other binding pocket for ATP^63,64^. To test whether the presence of WEE1 is required for ISR activation, RPE1 *TP53*^KO^ cells were depleted of WEE1, and subsequently treated with AZD1775. WEE1 depletion did not prevent ATF4 upregulation or GCN2 phosphorylation, indicating that ISR activation by AZD1775 does not require WEE1 presence (Fig. 7B). To address whether AZD1775 could similarly activate GCN2 kinase as previously described kinase inhibitors, *in vitro* kinase assays were performed with purified GCN2 kinase domain and the yeast eIF2α orthologue SUI2 as a substrate. Phosphorylation of eIF2α by GCN2 upon addition of the WEE1i AZD1775 was not observed (Suppl. Fig. 7A). Subsequently, activation of full-length human GCN2 was tested upon AZD1775 addition. GCN2 activation was measured in the presence of total cellular RNA, as purified uncharged tRNAs or ribosomes can stimulate the activity of GCN2 *in vitro*^65^. Moreover, tRNAs may be necessary for the paradoxical activation of GCN2 by ATP-competitive kinase inhibitors^64^. Again, phosphorylation of eIF2α by GCN2 was observed, which was not affected by addition of AZD1775 (Suppl. Fig. 7B). These observations suggest that AZD1775 does not directly trigger GCN2 kinase activity, in the absence of its coactivator GCN1 or ribosome components. Next, a NanoBRET assay^66^ was used to address the previously reported observation that AZD1775 could bind the recombinant kinase domain^67^. Specifically, we assessed the ability of the WEE1i AZD1775, Debio 0123 and ZNL 02-096, a WEE1 PROTAC consisting of AZD1775 linked to an E3 ubiquitin ligase, to displace a fluorescent tracer compound at the active site of luciferase-tagged WEE1 or GCN2 kinase domain in a cellular environment. In these assays, all three compounds potently displaced the fluorescent tracer from the WEE1 kinase domain at biologically relevant doses (Fig. 7C). Interestingly, no significant displacement at the GCN2 kinase domain was observed for AZD1775, Debio 0123 or ZNL 02-096 (Fig. 7D), at concentrations sufficient to activate GCN2 and induce ATF4 expression. In contrast, a non-specific pan-kinase inhibitor (CC1) did interact with the GCN2 kinase domain (Fig. 7D). Taken together, these data suggest that ISR activation by chemical WEE1i AZD1775, Debio 0123 or ZNL 02-096 is unlikely to be caused by direct interaction with the GCN2 kinase domain. Similar to the WEE1i, treatment of RPE1 *TP53*^KO^ cells with ZNL 02-096 resulted in upregulation of ATF4 expression and GCN2 phosphorylation (Suppl. Fig. 7C). Intriguingly, whereas AZD1775 treatment resulted in sustained ISR activity, we observed a decrease in ATF4 upregulation at 12 h after ZNL 02-096 treatment. Combined, these data show that ISR activation by chemically distinct WEE1i is independent of WEE1 presence, pointing at off-target activity of the WEE1i AZD1775, Debio 0123 and ZNL 02-096.

### WEE1i and PKMYT1i synergistically kill *TP53*^KO^ cells without WEE1i-mediated induction of the ISR

In line with WEE1 and PKMYT1 both regulating CDK1 activity, their combined inhibition showed synergy even at individual sub-cytotoxic doses^50,68^. Combined WEE1 and PKMYT1 inhibition in solid tumors is currently being evaluated in a phase 1/1b clinical trial (NCT04855656). To investigate whether WEE1i combined with RP6306, a PKMYT1i^69,70^, can synergistically induce cytotoxicity without WEE1i-mediated ISR activation, ATF4 protein expression was measured following treatment with a dose range of Debio 0123 (Fig. 8A, Suppl. Fig. 8A) or AZD1775 (Suppl. Fig. 8A, B). Debio 0123 treatment only resulted in ATF4 upregulation starting at 500 nM (Fig. 8A), whereas AZD1775-mediated ATF4 expression was observed at 20 nM onwards (Suppl. Fig. 8B).

**Figure 8.**
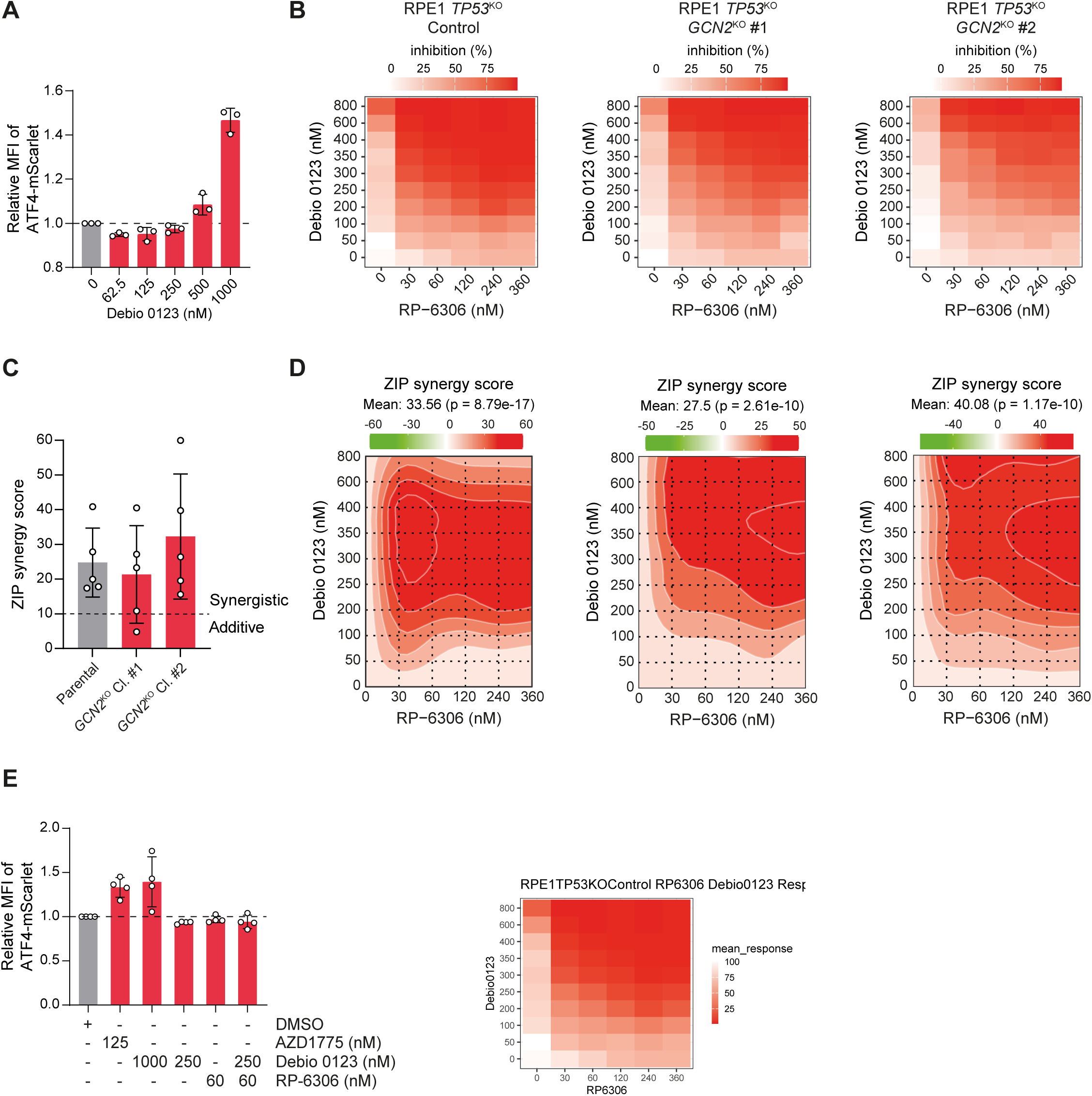
WEE1i and PKMYT1i synergistically induce cytotoxicity without WEE1i-induced ISR activation. **(A)** Relative ATF4-mScarlet MFI in RPE1 *TP53*^KO^ ATF4-mScarlet reporter cells after treatment with the indicated doses of Debio 0123. Data represent mean ± SD (n=3). **(B)** Cell viability in parental RPE1 *TP53*^KO^ *PAC*^KO^, *GCN2*^KO^ #1 and *GCN2*^KO^ #2 cells after treatment with Debio 0123 and RP-6306 at the indicated doses (n=5). **(C)** ZIP synergy scores from cell survival matrices shown in Figures 7B and 7D. Data represent mean ± SD (n=5). **(D)** Synergy plots with ZIP synergy scores of heatmaps shown in figure 7B. **(E)** Relative ATF4-mScarlet MFI in RPE1 *TP53*^KO^ ATF4-mScarlet reporter cells treated with the indicated doses of AZD1775, Debio 0123 and RP-6306. Data represent mean ± SD (n=4).

We observed synergy between WEE1i and PKMYT1i, with a stronger reduction in viability with the combination treatment compared to the highest doses of either single treatment (Fig. 8B, D; Suppl. Fig. C, E; Suppl. Table 2). Importantly, RPE1 *TP53*^KO^ cells treated with Debio 0123 and PKMYT1i showed a mean synergy (ZIP) score of 24.8, ranging from 17.4 and 40.9 between experiments (Fig. 8C). *GCN2*^KO^ cells required slightly higher doses of the combination treatment to reach the same viability reduction compared to parental cells (Fig. 8B, D), although the synergy between Debio 0123 and PKMYT1i treatment was maintained in *GCN2^KO^* cells, with mean ZIP scores of 21.3 and 32.3 for *GCN2*^KO^ #1 and #2, respectively (Fig. 8B-D). Importantly, synergy between Debio 0123 (250 nM) and RP-6306 (60 nM) was observed at doses that did not induce ATF4 expression, either in single agent treatment or in combination (Fig. 8A-E). Similar results were obtained with AZD1775 treatment in RPE1 *TP53*^KO^ cells, with mean ZIP scores of 22.4, ranging from approximately 9 to 33 (Suppl. Fig. 8C-E). Again, *GCN2*^KO^ cells required higher doses of both inhibitors to achieve the same level of toxicity as parental RPE1 *TP53*^KO^ cells, although *GCN2*^KO^ #1 and #2 cell lines still exhibited synergy with mean ZIP scores of 14.3 and 18.4 respectively (Suppl. Fig. 8C-E). Taken together, these findings indicate that combined treatment of low-dose Debio 0123 WEE1i and PKMYT1i induces cytotoxicity independently of ISR activation.

## Discussion

In this study, we report that the sensitivity of cells to clinically used WEE1i is determined by the integrated stress response (ISR) and ribosome quality control (RQC) pathways, in addition to its known effects on cell cycle perturbation. The WEE1i AZD1775 and Debio 0123 trigger activation of GCN2 in transformed and untransformed cells, in a cell cycle-independent manner. Specifically, loss of ISR-kinase GCN2, its binding partner GCN1, or its downstream effector ATF4 decrease WEE1i cytotoxicity. Yet, cells in which the ISR is inactivated remain susceptible to WEE1i-mediated cell cycle checkpoint override and synergistic effects of WEE1i with genotoxic chemotherapeutics and with PKMYT1 inhibition. Clearly, current WEE1i trigger two mechanistically distinct cytotoxic responses.

Remarkably, we found that activation of the ISR by the WEE1i used in this study was most likely independent of WEE1. A potential explanation for this observation is that the AZD1775 and Debio 0123 molecules directly bind and activate GCN2, analogous to other tyrosine kinase inhibitors^63,64,71^ that are known to bind the ATP-binding pocket of one of the monomers of the GCN2 dimer. In line with this hypothesis, previously reported kinome profiling studies employing phage display reported binding of AZD1775 to recombinant GCN2 kinase domain^67,72^. A more recent study also suggested this mode of action for AZD1775, when tested at high concentrations^73^. However, we did not find evidence for direct activation of GCN2 by WEE1i. Cellular NanoBRET assays showed neither AZD1775, Debio 0123 nor ZNL 02-096 binding to the GCN2 kinase domain at concentrations that cause activation of GCN2 and induction of ATF4 expression. Although it is technically possible that these WEE1i bind to another domain of GCN2 or only bind in the context of full-length GCN2, we currently favor the hypothesis that these agents trigger GCN2 activation via perturbation of translation. In support of this mechanism, i) GCN2 is not degraded by ZNL 02-096^72^, ii) *GCN1* inactivation confers resistance to AZD1775 treatment, and iii) ongoing translation is required for AZD1775 to trigger the ISR. That said, the exact cellular target of WEE1i that activates ISR remains elusive. Interestingly, the WEE1 PROTAC ZNL 02-096, which is based on AZD1775, only triggered transient GCN2 phosphorylation and ATF4 upregulation, arguing that the AZD1775 off-target molecule which is responsible for ISR activation is also degraded by ZNL 02-096.

The WEE1i-induced ISR activation resulted in an attenuation of global mRNA translation and upregulation of stress response factors. In the absence of eIF2A, GCN2 can still be activated, leading to eIF2α phosphorylation, which subsequently results in reduced cap-dependent translation and presumably increased translation of stress-related ISR target proteins. In a context where eIF2A is not available, translation of these stress-related ISR target proteins is likely reduced, possibly due to the requirement of eIF2A for their translation^32^, likely leading to critically low translation levels, which explains the increased sensitivity of *EIF2A^KO^* cells to WEE1i. Loss of ribosome rescue pathway factors also resulted in increased sensitivity to WEE1i treatment. This genetic context leads to increased ribosomal stress. Specifically, previous studies have shown that loss of *ZNF598* results in the readthrough of ribosomal stalling sites^58,74^. In yeast, loss of the ZNF598-ortholog Hel2 was demonstrated to increase GCN2 activation^75^. While in conditions of infrequent ribosome collisions ZNF598/Hel2 is activated, in conditions of excessive ribosome collisions, GCN2 will be activated as well^75^. Possibly, GCN2 activation in response to loss of RQC factors may become cytotoxic when GCN2 is additionally activated by WEE1i. In the setting that WEE1i activate GCN2, and RQC factor expression is lost, GCN2 might be occupied and therefore unable to respond to baseline collisions which are normally resolved by RQC factors. Therefore, WEE1i may lead to increased levels of stalled or collided ribosomes which are not resolved or recycled, subsequently resulting in an increased number of ribosomes per RNA containing a stall site. This model aligns with our socRNA data. In accordance with this, we also observed an increase in 80S ribosomal subunits after AZD1775 treatment, which could be due to reduced 80S splitting by ABCE1^76^.

In previous studies, WEE1i was studied as a single agent or in combination regimens with genotoxic agents to induce uncontrolled cell cycle progression in the presence of DNA lesions to trigger mitotic catastrophe^30^. Clinical evaluation of WEE1i demonstrated favorable responses along with considerable bone marrow toxicity^23,25,26,77^. Although ISR activation is a mechanism to restore homeostasis upon stress, chronic stimulation of ISR kinases can induce apoptosis^78^. Therefore, ISR activation may contribute to these adverse side-effects. Indeed, we observed that ISR activation by WEE1i also occurred in non-transformed blood cells. However, since unstimulated blood cells have low levels of mRNA translation, we anticipate that ISR activation will only manifest itself in subpopulations that are metabolically active.

The notion that WEE1i in tumor cells exert two mechanistically distinct effects – loss of cell cycle control and ISR activation – may challenge biomarker identification for WEE1i eligible patients. Indeed, biomarker studies have predominantly focused on tumor characteristics related to defective cell cycle control (i.e. *TP53* inactivation) or replication stress (i.e. *SETD2* loss, *CCNE1* amplification). To improve patient benefit, novel WEE1i could be developed, that do not activate the ISR. Alternatively, combination strategies based on combining low doses of currently used WEE1i with other agents could also be considered. Indeed, our data show that the previously described synergistic combination of the WEE1i Debio 0123 with PKMYT1i^68^ is effective at doses that do not trigger ISR activation. Combination of Debio 0123 with RP-6306 at low doses may therefore represent a therapeutic strategy to limit off-target cytotoxicity through ISR activation, while maintaining efficacy against cancer cells. Similarly, the anticipated development of next-generation dual PKMYT1/WEE1 inhibitors may provide therapeutic advantages of CDK1/2 modulation without the potential liability of ISR induction. However, it is also possible that ISR induction may be therapeutically desirable under some circumstance. Indeed, based on our genetic screening results, tumors with defective RQC pathways or loss of eIF2A are expected to be increasingly sensitive to current WEE1i.

## MATERIALS AND METHODS

### Cell line models

RPE1 hTERT *TP53*^KO^ (RPE1 *TP53*^KO^) cells were cultured in Dulbecco′s Modified Eagle′s Medium (Gibco) with low glucose (1 g/L; DMEM-L), supplemented with 10% fetal calf serum (FCS, Lonza) and 50 units/mL penicillin and 50 μg/mL streptomycin (Gibco, 15140122). RPE1 hTERT *TP53*^KO^ (RPE1 *TP53*^KO^) *PAC*^KO^ cells, RPE1-FLAG-Cas, HEK293T cells, and derivative lines were maintained in DMEM with 4.5 g/L glucose (DMEM-H), 1 mM sodium pyruvate, supplemented with 10% fetal calf serum (FCS). Media for RPE1 *TP53*^KO^ *ZNF598*^KO^, *ATF4*^KO^, and *GCN2*^KO^ cells were supplemented with MEM non-essential amino acids, or DMEM with 1 g/L glucose supplemented with 10% FCS and 50 units/mL penicillin and 50 μg/mL streptomycin. For CRISPR/Cas9 screens, RPE1 and HeLa cells were grown in DMEM-H, 1 mM sodium pyruvate, 1x penicillin-streptomycin (Wisent), and 10% FCS. HAP1 cells were cultured in Iscove’s Modified Dulbecco’s Medium (IMDM, Gibco), supplemented with 2 mM L-glutamine (Gibco), 50 units/mL penicillin, 50 μg/mL streptomycin (Gibco, 15140122) and 10% FCS. OVCAR3 cells were cultured in Roswell Park Memorial Institute (RPMI)-1640 media supplemented with 50 units/mL penicillin, 50 μg/mL streptomycin, and 20% FCS. Peripheral Blood Mononuclear Cells (PBMCs) were cultured in RPMI-1640 medium supplemented with 10% FCS and 2 mM L-glutamine. HCC38 cells were maintained in RPMI-1640 medium supplemented with 50 units/mL penicillin, 50 μg/mL streptomycin, and 10% FCS. U2OS cells stably expressing tetR, ALFA-Nb-CAAX, and STAb-GFP were cultured in DMEM-H with 50 units/mL penicillin, 50 μg/mL streptomycin, and 5% FCS. All cells were cultured at 37°C and 5% CO_2_.

### Inhibitors and fine chemicals

The chemicals used in this study were AZD1775 (#1494, Axon Medchem or S1525, Selleck Chemicals), Debio 0123 (S9778, Selleckchem), A92 (#2720, Axon Medchem), RP-6306 (gift from Repare Therapeutics), AZD7762 (#1399, Axon Medchem), VE-822 (#2452, Axon Medchem), ZNL 02-096 (7240, Tocris), thapsigargin (T9033, Merck), doxorubicin (sc-280681, Santa Cruz Biotechnology), nocodazole (M1404, Sigma), palbociclib (1505, Axon Medchem), integrated stress response inhibitor (ISRIB; SML0843, Sigma or S7400, Selleck Chemicals), salubrinal (S2923, Selleck), and Emetine (E521535, Toronto Research Chemicals).

### Haploid genetic screen

HAP1 cells were mutagenized as described previously^30,79,80^. In summary, ∼6 × 10^7^ cells were transduced with pGT gene-trap retrovirus. Mutagenized cells were treated for 9 days with AZD1775 (125 nM) prior to fixation in ‘Fixation Buffer 1’ (Becton Dickinson). Genomic DNA was isolated from ∼4 × 10^7^ cells. DNA was subjected to a linear amplification (LAM)-PCR protocol, and sequenced using the Genome Analyzer platform (Illumina). Gene trap insertion mapping and data analysis was performed as described previously^81^. In brief, the dataset was normalized to combined insertion counts of four HAP1 wildtype datasets^79^. A binomial test was performed for the distribution of sense and antisense orientation insertions in one AZD1775-treated HAP1 dataset. Loss of a gene was considered sensitizing to AZD1775 when the sense-over-antisense insertion ratio was decreased in the AZD1775-treated HAP1 dataset compared to the four HAP1 wildtype datasets^79,82^ (SRA accession no. SRP058962). To this end, two-sided Fisher’s exact tests were performed for each gene against all wildtype datasets, and the gene was considered a sensitizer when it passed all Fisher’s tests (p < 0.05; effect size 20%). Sequencing data have been deposited at the BioProject Archive of the NCBI under accession PRJNA1152540.

For pathway analysis, sense insertion ratios and insertion frequency of WEE1i-treated cells were compared with 4 DMSO-treated control datasets. P values for each gene were log-10 transformed, and genes were ranked according to average P value. The top 250 genes of which WEE1i-treated cells had increased more antisense insertions were analyzed by DAVID^82^, using the KEGG pathway gene sets. Only gene sets with a false-discovery rate <0.05 are indicated.

### Genome-wide CRISPR screens with Toronto KnockOut sgRNA libraries

For CRISPR/Cas9 screens, RPE1 *TP53*^KO^ Flag-Cas9 cells were transduced at low multiplicity of infection (MOI 0.3-0.5) with Toronto KnockOut (TKO) v1 90k sgRNA or v2 71k sgRNA libraries for the sensitizer and resistance screen, respectively^83–85^. The next day, infected cells were selected with 20 µg/ml puromycin with trypsinization 24 h after selection and then expanded in the presence of puromycin and blasticidin. Cells were then split into technical duplicates maintaining the appropriate library coverage on day T0. Cells were split once more (day T3) before addition of AZD1775 on day T6. For the negative selection screen, cells were split every three days in the presence of 80 nM AZD1775. Screens were stopped at an estimated 10 population doublings (T21 for AZD1775 comparisons shown in this study). For the positive selection screen with AZD1775, RPE1 *TP53*^KO^ Flag-Cas9 cells were split once dishes were 90% confluent (6-9 days after first addition of 200 nM AZD1775) and then collected a second time (T18-T21). Genomic DNA was extracted from samples taken at each time point (T0, T6, T18-21) using the QIAamp DNA Blood kit (Qiagen). Genomic DNA was normalized by measurement on a Nanodrop spectrophotometer. TKO sgRNA cassettes were isolated by PCR using KAPA HiFi HotStart ReadyMix (Roche). Illumina i5 and i7 sequencing multiplex barcodes were added in a second round of PCR and libraries were purified by agarose gel electrophoresis. Sequencing libraries were processed on an Illumina NextSeq500 platform at the Lunenfeld-Tanenbaum Research Institute NBCC facility (https://nbcc.lunenfeld.ca). sgRNA representation was determined by aligning sequencing reads to reference libraries and then using read counts as input for MAGeCK or DrugZ analysis to compute gene-level sgRNA enrichment and depletion, respectively. DrugZ values are provided in Suppl. Table 4. Negative log10-transformed positive MAGeCK scores for individual genes were plotted. Raw MAGeCK scores are provided in Suppl. Table 5.

### CRISPR/Cas9-based gene editing

A sgRNA targeting exon 4 of *EIF2A* (Suppl. Table 3) was cloned into the PX458 vector, which was a gift from Fengh Zhang (Addgene #48138). Plasmids were introduced in RPE1 *TP53*^KO^ and OVCAR3 cell lines using Fugene HD and were flow-sorted based on GFP expression.

Genomic DNA was isolated using QuickExtract DNA solution (Lucigen), followed by PCR using Taq polymerase (New England Biolabs) according to manufacturer’s instructions. Samples were sequenced by Eurofins Genomics using a Mix2Seq kit (Eurofins). TIDE was used to identify the mutational pattern after CRISPR-Cas9 mediated editing of *EIF2A*^86^. RPE1 *TP53*^KO^ *EIF2A*^KO^ cells had 93.8% out-of-frame mutations. OVCAR3 *EIF2A*^KO^ #1 cells had 36% out-of-frame mutations, and OVCAR3 *EIF2A*^KO^ #1 cells had 32.3% out-of-frame mutations and 13.4% in-frame indels.

RPE1 *TP53*^KO^ *PAC*^KO^ *ATF4*^KO^ and *GCN2*^KO^ CRISPR/Cas9 edited cells were generated by transfection of parental cells with Cas9 ribonucleoprotein complexes (RNP) using Lipofectamine CRISPRMAX reagent (Life Technologies) and single clones were selected. sgRNAs were generated by *in vitro* transcription using the TranscriptAid T7 High Yield kit (ThermoFisher) (Suppl. Table 3). RPE1 *TP53*^KO^ *PAC*^KO^ *ZNF598*^KO^ cells were generated by transfecting parental cells with plentiCRISPRv2 derived sgRNA expression vector (plentiGuide; Suppl. Table 3) and selecting for single clones. To assess sgRNA editing efficiency in polyclonal populations, transfected cells were passaged at least once before collection for genomic DNA extraction, PCR, and Sanger sequencing. Cas9 editing was assessed using the tracking of indels by decomposition (TIDE, Netherlands Cancer Institute) or inference of CRISPR edits (ICE) algorithm (Synthego). CRISPR editing in clonal cell lines was confirmed by Sanger sequencing and immunoblotting.

### DNA cloning and retroviral transduction

The ATF4-mScarlet-NLS insert of pLHCX-ATF4 mScarlet NLS, which was a gift from David Andrews (Addgene, #115970) was obtained by PCR (Suppl. Table 3). Restriction digest was applied on pMSCV-blast (Addgene, #75085) using SalI (ThermoFisher, FD0644) and BamHI (ThermoFisher, FD0054) according to the manufacturer’s instructions. The ATF4-mScarlet NLS insert was cloned into the digested pMSCV-blast using NEBuilder HIFI DNA assembly Master Mix (New England Biolabs) according to the manufacturer’s instructions. Selected colonies were sequenced by Eurofins Genomics using a Mix2Seq kit (Eurofins). TIDE analysis was used to check the integration of the insert^86^.

The pMSCV-ATF4-mScarlet-NLS construct was transfected into HEK293T cells along with pRetro-VSV-G and pRetro-gag/pol, and subsequently, RPE1 *TP53^KO^* cells were transduced with retrovirus. Successfully transduced cells were selected with 7.5 µg/ml blasticidin S HCl (ThermoFisher, R21001) and sorted monoclonally using the Sony SH800S cell sorter.

### siRNA transfection

For knockdown of *WEE1*, RPE1 *TP53^KO^* cells were transfected with siRNA targeting *WEE1* (ThermoFisher, HSS111337) using oligofectamine (Invitrogen, 12252-011) according to the manufacturer’s instructions. After 72 h of transfection, cells were treated with AZD1775 for the indicated time points.

### Viability assays

MTT assays: For assessing the sensitivity of *EIF2A^KO^*cells to AZD1775, RPE1 *TP53*^KO^ cells were seeded at a density of 200 cells/well of a 96-well plate, while OVCAR3 cells were seeded at 500 cells/well. For AZD1775 and cisplatin synergy assays, 100 cells/well were seeded for RPE1 *TP53^KO^ PAC*^KO^ control cells, and *GCN2^KO^* cells were seeded at a 200 cells/well density. For AZD1775 and cisplatin synergy assays treated with A92, RPE1 *TP53^KO^* cells were seeded at a density of 200 cells/well. For PKMYT1i and WEE1i synergy assays, RPE1 *TP53^KO^* PAC^KO^ control and *GCN2*^KO^ cells were plated at a density of 100-400 cells/well. 24 h after seeding, drugs were added at the indicated doses for 5 days. 5 mg/mL of methyl-thiazol tetrazolium (MTT; Sigma, M2128) was added for 4 h, after which medium was removed and formazan crystals were dissolved in DMSO. Absorbance values were determined using a spectrophotometer at a wavelength of 520 nm. For synergy analysis, SynergyFinder version 3.0 (synergyfinder.fimm.fi) was used to generate heatmaps and calculate ZIP synergy scores^87^.

Clonogenic assays: RPE1 *TP53*^KO^ cells were seeded at a density of 200 cells/well of a 6-well plate, while OVCAR3 cells were seeded at 500 cells/well. 24 h after seeding, drugs were added at the indicated doses for 10 days. Culture medium was removed, and cells were fixed and stained using 0.2% Coomassie Brilliant Blue solution containing 50% methanol (Merck) and 14% acetic acid (Merck) in PBS. Plates were washed in tap water and air-dried overnight.

Images of the plates were loaded into Adobe Photoshop, and a concentric circle was drawn to isolate the same area for each well. The magic wand tool was used to select colonies, and data per colony was exported. Subsequently, data for each colony was loaded into RStudio, and all areas with a circularity measure below 0.1 and an area below 12, which represent technical artifacts, were removed. The experiment was summarized to calculate area covered per well.

For experiments performed with RPE1 *TP53*^KO^ *PAC*^KO^ *ATF4*^KO^ and *GCN2*^KO^ clones, cells were seeded at a density of 100 cells/well in 6-well plates or at 300-1000 cells/dish in 10 cm dishes containing dilutions of AZD1775 and vehicle. After 12-20 days, surviving colonies were fixed and stained with crystal violet (0.4% w/v crystal violet (Sigma), 20% methanol). Quantification of experiments was performed using a GelCount scanner (Oxford Optronix) and surviving fractions were calculated by normalization to vehicle treatments.

For experiments with the IncuCyte S3 Live-Cell Analysis Instrument (Essen Bioscience, Sartorius), RPE1 *TP53*^KO^ *PAC*^KO^ cells were seeded in 96-well plates (100-300 cells per well) in the presence of serial dilutions of the indicated concentrations of AZD1775 and vehicle controls. Cells were imaged with a 10X objective once the vehicle controls reached confluency (approximately 6 to 9 days). Cell confluency was calculated automatically using IncuCyte segmentation algorithms based on phase contrast images and normalized to vehicle treatments.

For fluorescence-based cell competition assays, RPE1 *TP53*^KO^ FLAG-Cas9 expressing cells were transduced at high MOI (MOI: 1) with plentiCRISPRv2 derived sgRNA expression vectors (plentiGuide) encoding EGFP-NLS or mCherry-NLS fluorescent proteins. For mCherry control expression vectors, β-galactosidase (*LacZ*) targeting sequences were used. Two days after infection (T_-1_), cells were seeded at low density in a 1:1 ratio in 12-well or 24-well plates to begin imaging the next day (T0) on a GE InCell Analyzer 6000 confocal microscope using a 4x objective. Where indicated, drug treatments were initiated on T_1_ and cultures were split in 1:20 fractions each time cells reached 90% confluency (approximately 3 to 4 days of growth). Cell fitness was calculated in each experimental condition by charting the ratio of GFP-positive to mCherry-positive cells at each imaging timepoint and normalizing the GFP:RFP ratio on day T_0_ to 1. Therefore, defects in cell fitness are presented as values < 1 and gain of fitness is represented by values > 1. The total number of GFP and mCherry-positive nuclei were segmented and counted in each well of an assay plate using a custom Acapella script (PerkinElmer).

### Western blotting

Cells were trypsinized and lysed in Mammalian Protein Extraction Reagent (M-PER™, ThermoFisher) supplemented with Halt phosphatase inhibitor and Halt protease inhibitor (Thermo Scientific). Protein concentration was measured using Pierce BCA Protein Assay Kit (Thermo Scientific). Proteins were separated by SDS-PAGE gel electrophoresis and transferred to methanol-activated PVDF membranes (Immobilon) using the Trans-Blot Turbo Transfer System. Subsequently, membranes were blocked in 3% Bovine Serum Albumin Fraction V (Sigma-Aldrich) or 5% skim milk (Sigma-Aldrich) in 0.05% TBS-Tween (TBST). Immunodetection was performed using antibodies directed against puromycin (MABE343, Merck, 1:10000), GCN2 (3302, Cell Signaling, 1:1000), pGCN2 (ab75836, Abcam, 1:500), ATF4 (11815, Cell Signaling, 1:1000), Vinculin (ab129002, Abcam, 1:2000), HSP90 (sc-13119, Santa Cruz, 1:5000), pCDK1/2/3/5 (ab133463, Abcam, 1:1000), eIF2A (ab169528, Abcam, 1:1000), WEE1 (4936S, Cell signaling, 1:1000), 1:1000 rabbit anti-cyclin B1 (Santa Cruz, sc-752), followed by staining with secondary antibodies Goat Anti-Rabbit Immunoglobulins/HRP (Dako, P0448) or Rabbit Anti-Mouse Immunoglobulins/HRP (Dako, P0260). Chemiluminescence detection was performed using Lumi-Light (Lumi-Light, Roche Diagnostics) or SuperSignal West Femto (SuperSignal, ThermoFisher) using ChemiDoc XRS+ System (Bio-Rad).

For the RPE1 *TP53*^KO^ *PAC*^KO^, *GCN2*^KO^, *ATF4*^KO^, ZNL 02-096 and emetine immunoblot experiments, whole cell lysates were prepared by scraping cells in warm 2x Laemmli sample buffer (2% SDS, 167 mM Tris HCl pH 6.8, 20% glycerol, 0.02% bromophenol blue, 2 mM DTT). Proteins were separated by SDS-PAGE and the gel was stained with Ponceau S. Afterwards, proteins were transferred onto nitrocellulose membranes and subsequently blocked with 5% milk or in 0.1% TBST and blotted with the following primary antibodies for 1 h at room temperature (RT) or overnight at 4°C: 1:1000 rabbit anti-phospho-eIF2a Ser51 (Cell Signaling, #3398), 1:1000 rabbit anti-phospho-eIF2a Ser51 (Cell Signaling, #9721), 1:1000 rabbit anti-eIF2a (Cell Signaling, #9722), 1:1000 rabbit anti-phospho-GCN2 Thr899 (Abcam, ab75836), 1:1000 rabbit anti-GCN2 (Abcam, ab134053), 1:1000 rabbit anti-ZNF598 (Sigma, HPA041760), 1:1000 rabbit anti-PERK (Cell Signaling, #5683), 1:1000 rabbit anti-phospho-cdc2 (CDK1) Tyr15 (Cell Signaling, #4539), 1:500 rabbit anti-ATF4 (Cell Signaling, #11815), 1:500 mouse anti-ATF4 (Cell Signaling, #97038), 1:500 rabbit anti-WEE1 (Cell Signaling, #13084). Blots were stained with the following secondary antibodies (1:10000 in 5% milk in 0.1% TBST) for 1 h at RT and chemiluminescent signal (SuperSignal West Dura, Thermo Fisher) was captured on film: horseradish peroxidase (HRP) conjugated AffiniPure goat anti-rabbit IgG (Jackson ImmunoResearch 111-035-144), HRP AffiniPure goat anti-mouse IgG (Jackson 115-035-003), HRP sheep anti-mouse IgG (Amersham NA931V).

### Puromycin assays

To assess protein synthesis rates, 20 μg/mL of puromycin was added to cells for 10 min before harvesting and lysis to label nascent proteins with puromycin. Further sample processing was the same as the general western blotting procedure as mentioned above. Quantification of puromycin incorporation was performed by measuring grey values of puromycin lanes, loading control bands and background in Adobe Photoshop. Grey values were reverted, background signal values for puromycin and loading controls were subtracted from their respective sample values. Loading control band values were subtracted from puromycin band values. Treated conditions were normalized to the control condition.

### Metabolic labeling with L-azidohomoalanine

To assess bulk mRNA translation, RPE1 *TP53*^KO^ *PAC*^KO^ cells were seeded in 15 cm dishes and then treated or not with AZD1775 and ISRIB for 24 h. Cells were washed once in PBS and then grown in DMEM high glucose without glutamine, methionine, or cysteine (Wisent) with 10% FBS containing 25 μM Click-iT L-azidohomoalanine (AHA, Thermo Fisher) for 2 h. Cells were collected by trypsinization and washed twice with PBS before lysis in buffer containing 1% SDS, 50 mM Tris pH 8.0, 1x complete protease inhibitor cocktail (Roche), and 150 U/ml benzonase nuclease (Sigma). Whole cell lysates were incubated on ice for 30 min and pelleted by centrifugation in a refrigerated microcentrifuge for 10 min at 4°C. Supernatants were normalized using a Nanodrop spectrophotometer and 1 μl of 10 mM IRDye800CW-DBCO (Li-Cor Bosciences) was added to each sample and incubated for 2 h at RT in the dark. Excess dye was removed by centrifugation of samples through Zeba spin desalting columns (7K MWCO, 0.5ml, Thermo Fisher) according to the manufacturer’s instructions. Reactions were boiled in 1X Laemmli sample buffer and SDS-PAGE was performed with Novex 4-12% Tris-Glycine gels (Invitrogen). IRDye800CW infrared fluorescence was detected on an Odyssey 9120 imager (Li-Cor) and total proteins were stained with Coomassie staining solution (0.1% w/v Coomassie Brilliant Blue, 20% methanol, 10% acetic acid). Total AHA labeled proteins were quantified by densitometry in Fiji (ImageJ) with background IRDye800CW fluorescence subtracted from measurements and normalized to AZD1775 and ISRIB vehicle (DMSO).

### Flow cytometry

For cell cycle analysis, RPE1 *TP53*^KO^ *PAC*^KO^ control, *ATF4*^KO^ #1, *ATF4*^KO^ #2, *GCN2*^KO^ #1, and *GCN2*^KO^ #2 cells were seeded at a density of 1.20-1.75*10^5^ cells/T25 and treated with 500 nM doxorubicin for 1 h before the addition of 62.5 ng/ml nocodazole to arrest cells in mitosis. Additionally, in RPE1 *TP53^KO^*-only experiments, cells were treated with A92 and AZD1775 simultaneously with 500 nM doxorubicin. After 16 h of treatment, cells were harvested and fixed in ice-cold 70% ethanol. Cells were stained with primary antibodies against phospho-Ser/Thr-Pro MPM-2 (05-368, Millipore, 1:100) and γH2AX (#9718, Cell Signaling, 1:100), and Alexa-Fluor 488 and 647 conjugated secondary antibodies (A32728 and A21206, ThermoFisher). DNA was stained using Propidium Iodide (PI; Sigma, P4864) while lysing RNA using RNAse A (Sigma, R6513). Samples were measured on a Quanteon NovoCyte (NovoCyte Quanteon, Agilent) and analyzed using FlowJo VX.

For ATF4-mScarlet reporter experiments, RPE1 *TP53^KO^*ATF4-mScarlet-NLS cells were seeded at a density of 30,000 cells/well in a 6-well plate. Cells were treated with indicated doses of thapsigargin, AZD1775, Debio 0123, A92, RP-6306, AZD7762 or VE-822. After 24 h, cells were measured on a Quanteon flow cytometer (NovoCyte Quanteon, Agilent) and analyzed using FlowJo VX. For the G1 arrest experiments, cells were plated at 90,000 cells/T25 density and treated with 62.5 ng/ml nocodazole or vehicle for 16 h before performing mitotic shakeoff, or in case of nocodazole-free controls, harvested by trypsinization. Subsequently, nocodazole was washed out and the cells were replated in the presence of 1.25 µM palbociclib or vehicle. After the cells had adhered, 1 µM AZD1775 and 1 µM A92 were added for approximately 20 h. After treatment, cells were harvested, measured on a Quanteon NovoCyte (NovoCyte Quanteon, Agilent), and analyzed using FlowJo VX.

### PBMC activation

PBMCs were seeded at 1.25M cells/ml density in 12-well plates. CD3^+^ cells were activated with a ratio of 1:8 Dynabeads Human T-Activator CD3/CD28 beads (Gibco, 11131) per cell for 4 days before the addition of inhibitors. 24 h after the addition of AZD1775 and thapsigargin, the PBMCs were pelleted and subsequently prepared for western blot as described.

### DNA Fiber assays

RPE1 *TP53*^KO^ and OVCAR3 cells were seeded at 50,000 cells/well in a 6-well plate and incubated for 24 h. Cells were pulse-labeled with 25 μM 5-iodo-2’-deoxyuridine (IdU; Sigma-Aldrich) for 20 min, washed three times with pre-warmed culture media, and subsequently pulse-labeled with 250 μM 5-chloro-2’-deoxyuridine (CldU; Sigma-Aldrich) for 20 min. Cells were trypsinized and diluted to a concentration of 50,000 cells/mL. Cells were lysed in lysis buffer (0.5% sodium dodecyl sulfate, 200 mM Tris pH 7.4, 50 mM EDTA) and allowed to spread by gravity flow at an angle of approximately 20 degrees. Slides were air-dried, fixed in methanol/acetic acid (3:1 ratio) for 15 min, and denatured in 2.5 M HCl for 1.5 h. A primary antibody directed against BrdU (BD Biosciences, 347580, 1:250, mouse) was used to detect IdU, while CldU was detected using a primary antibody directed against BrdU (Abcam, ab6326, 1:1000, rat). Slides were incubated with primary antibodies for 2 h at RT, followed by incubation with Alexa Fluor 488- or 647-conjugated secondary antibodies (1:500) for 1.5 h. Images were acquired on a Leica DM-6000 fluorescence microscope, equipped with Leica Application Suite software. The lengths of IdU and CldU tracks were measured using ImageJ.

### Immunofluorescence microscopy

18 mm square coverslips were transferred into 6-well plates and sterilized in a microwave at 900W for 2 min. RPE1 *TP53*^KO^ and *EIF2A*^KO^ cells were seeded onto the coverslips at a 30,000 cells/well density and incubated for 24 h. Soluble proteins were extracted by incubation with CSK buffer pH 7 (10 mM PIPES pH 7, 100 mM NaCl, 300 mM sucrose, 3 mM MgCl_2_, 0.5% Triton X-100) for 2 min on ice before fixation using 2% paraformaldehyde, permeabilization using PBS-0.1% Triton X-100 and blocking with 3% BSA V. Cover slips were incubated overnight at 4°C in primary antibodies directed against γH2AX (05-636, Millipore, 1:300). Cells were then incubated with an Alexa Fluor 647-conjugated secondary antibody (A32728, ThermoFisher) for 2 h at RT (1:500) and stained with DAPI. Finally, cells were mounted with Vectashield anti-fade mounting medium (Vectorlabs). Imaging was performed using the Zeiss Axio Imager Z2 and analysis was performed in FIJI. Experimental details for quantitative image-based cytometry (QIBC) of ATF4 levels are provided in the Supplementary Methods.

### SILAC labeling

Labeled isotopes (10Arg and 8Lys), SILAC DMEM (5 g/L glucose), and SILAC FBS were obtained from Silantes. Medium supplemented with labeled isotopes is called “heavy”, while medium without labeled isotopes is called “light”. RPE1 *TP53*^KO^ and RPE1 *TP53*^KO^ *EIF2A*^KO^ cells were cultured in either heavy or light medium for 5 passages (19 days) to saturate all proteins with isotope-labeled or unlabeled amino acids. Cells were treated with DMSO or 125 nM of AZD1775 to avoid excessive cell death in the more sensitive *EIF2A*^KO^ cell line. For all samples, a replicate was performed swapping the heavy and light label to obtain n=2 within the experiment. At 0, 6, 24, and 48 h, samples were taken by trypsinization, washing with PBS, and the dry cell pellet was snap-frozen and stored at -20°C. SILAC samples were lysed in MPER supplemented with protease and phosphatase inhibitors, and protein concentration was determined using a BCA assay. Heavy and light samples were mixed to obtain a 1 to 1 ratio of proteins containing 50 μg of protein in 50 μL of MPER. Per sample, 6X loading buffer with 10% β-mercaptoethanol was added and samples were boiled for 5 min at 95°C. Samples were loaded onto an acrylamide gel, after which proteins were digested in gel with trypsin. Digested peptides were analyzed using a linear ion trap-Orbitrap hybrid mass spectrometer (LTQ-Orbitrap, ThermoScientific). The MS data were analyzed with MaxQuant and mapped on the UniProt human proteome build with accession number UP000005640. Protein ratios were calculated from normalized protein counts and converted to a log fold change.

### GCN2 kinase assays

To probe GCN2 (EIF2AK4) kinase activity *in vitro*, 20 ng of truncated GCN2 kinase domain (KD, residues 192-1024, Sigma SRP5218) or full-length recombinant GCN2 expressed and purified from HEK293T cells was mixed with 1 μg of purified eIF2a from *S. cerevisiae* (SUI2) in kinase assay buffer (50 mM HEPES pH 7.4, 100 mM KOAc, 5 mM MgAc2, 6.25 mM b-glycerophosphate, 18.75 mM MgCl2, 1 mg/ml BSA, 1 mM DTT) on ice. For reactions with full-length GCN2, microtubes were incubated with 10 mg/ml BSA overnight, then washed twice with dH2O before use. Reactions were started by adding ATP mix (to a final concentration of 0.4 mM ATP) and transferring microtubes to a 32°C heat block. Samples were taken at 0, 5, and 10 min, then mixed with equal volumes of 2X Laemmli sample buffer for SDS-PAGE and probed with the indicated antibodies. AZD1775 or vehicle (DMSO) was titrated into reactions at the indicated final concentrations. For reactions with full length GCN2, total RNA (100 ng from HEK293T cells) was supplemented to reactions where indicated.

### NanoBRET assays

To assess AZD1775, Debio 0123 and ZNL 02-096 binding to GCN2 kinase domain in cells, HEK293T cells were reverse transfected with NanoLuc-EIF2AK4 (Domain 2) (Promega, NV3051) or WEE1-NanoLuc (Promega, NV2231) and transfection carrier DNA in opaque 96-well plates (Corning, #3917). 24 h after transfection, fluorescent tracer K-10 (NanoBRET TE intracellular kinase assay K-10, Promega, N2640) was added to cells at a final concentration of 0.5 μM and plates were mixed for 15 seconds on an orbital shaker at 900 rpm. Serial dilutions of AZD1775 or CC1 pan-kinase inhibitor (Promega, N2261) were added to cells and plates were incubated for 3 h. NanoBRET signal was measured by incubating cells with diluted Nano-Glo substrate according to the manufacturer’s instructions and detecting donor and acceptor emissions with an EnVision microplate reader (PerkinElmer). BRET ratios corresponding to tracer occupancy were determined by dividing acceptor emission by donor emission and multiplying by 1000 to compute mBu units. Background mBu signal was measured in wells where tracer compound was absent. Fractional occupancy of the GCN2 kinase domain for AZD1775, Debio 0123, ZNL 02-096 and CC1 was calculated using the following formula: [1 – (X-Z)/(Y-Z)] * 100, where X = mBu in the presence of test compound and tracer, Y = mBu in the presence of vehicle (100% BRET signal), and Z = mBu in the absence of tracer compound (no BRET control).

### Single cell EdU sequencing (scEdU-seq)

To assess replication dynamics, scEdU-seq was performed as described previously^88^. Briefly, RPE1 *TP53^KO^* control and *EIF2A^KO^* cells were treated as indicated and pulsed with 10 µM EdU for 15 min. The cells were subsequently harvested and fixed. Subsequently, pelleted cells were subjected to an EdU click reaction using to EdU-Click 647 imaging kit (Invitrogen) using the manufacturer’s instructions with minor modifications. Instead of Azide-647, 2 mM of azide-PEG3-biotin conjugate (Sigma) was used and supplemented with 6 mM tris((1-hydroxy-propyl-1H-1,2,3-triazol-4-yl)methyl)amine (Jena Bioscience). After this, cells were sorted to achieve single cells per well and lysed with Proteinase K. To digest the genome, NIaIII was added, and the resulting overhangs were processed to blunt ends using Klenow large fragment. End-repaired DNA fragments were A-tailed, and these fragments were ligated to T7 promoter containing adapters with cell barcodes and UMI. Further details of cDNA preparation are provided in the Supplementary Methods. cDNA libraries were sequenced using v2.5 chemistry on a NextSeq500 or NextSeq2000 (Illumina; NextSeq control software v.2.2.0.4; RTA v.2.4.11) with 100 cycles for read 1 (cell index and tables were filtered for the presence of a NlaIII restriction site, a mapping quality >30, the molecule has a pair of reads assigned, the molecule is unique and should not have alternative alignment positions in the genome. The code for analysis and plotting is available on GitHub: https://github.com/jervdberg/WEE1

### Polysome profiling

To assess bulk mRNA translation, RPE1 *TP53*^KO^ *PAC*^KO^ cells were treated with 250 nM AZD1775 or DMSO for 6 h. Cells from six 15 cm dishes per condition were pre-treated with 100 μg/ml cycloheximide (CHX) for 5 min and washed once with cold PBS containing 100 μg/ml CHX. Cells were scraped in PBS-CHX on ice and pelleted before resuspension in hypotonic buffer (5 mM Tris HCl pH 7.5, 2.5 mM MgCl2, 1.5 mM KCl, 1 mM DTT, 100 U/ml RNasin recombinant RNase inhibitor (Promega), 100 μg/ml CHX). Cells were lysed by addition of 10% Triton X-100 (final concentration 0.5%) and 10% sodium deoxycholate (0.5% final) and vortexing for 5 seconds. Whole cell lysates were cleared by centrifugation for 10 min at maximum speed in a cooled tabletop microcentrifuge. Lysates were normalized by measuring absorbance at 260nm (A260) on a Nanodrop spectrophotometer. For each sample, 15 A260 optical units were loaded on 5-50% sucrose gradients prepared in polysome buffer and centrifuged in an Optima XPN-80 ultracentrifuge with SW41Ti rotor (Beckman Coulter) at 39000 rpm for 2 h at 4°C. Polysome profiles were collected with a density gradient fractionation system (Brandel) measuring absorbance at 254nm.

### Stopless ORF circular (soc)RNA assays

To investigate changes in ribosome kinetics, assays using socRNAs were performed as described previously^59^. Briefly, U2OS cells expressing STAb-GFP, tet repressor and ALFA nanobody-CAAX were seeded at 25% confluency in 96-well glass-bottom plates (Matriplates, Brooks Life Science Systems) and after 24 h, were transfected with the plasmid encoding the socRNA using Fugene (Promega). Subsequently, cells were treated with 300 nM AZD1775 or vehicle for 24 h. Imaging was done the following day by replacing the medium with pre-warmed imaging medium (CO_2_-independent Leibovitz’s-15 medium (Gibco) containing 5% FCS (Sigma-Aldrich) and 1% penicillin/streptomycin (Gibco)). 90 min prior to the start of imaging, doxycycline (Dox, 1 µg/mL) was added to the cells to induce socRNA expression. All live-cell imaging experiments were performed at 37°C using the Nikon TI inverted microscope with NIS Element Software equipped with a perfect focus system, a Yokagawa CSU-X1 spinning disc, an iXon Ultra 897 EM-CCD camera (Andor), and a motorized piezo stage (Nanocan SP400, Prior). The microscope was equipped with a temperature-controlled box. A 100x 1.49 NA oil immersion objective was used for all imaging experiments. To precisely determine the number of translating ribosomes on socRNAs, the translation inhibitor puromycin (0.1 mg/mL; ThermoFischer Scientific) was added to cells 1-2 h after the start of imaging to induce nascent chain release. For live-cell imaging of socRNAs, the x, y positions for imaging were chosen based on the presence of translating socRNAs in cells. Images were acquired every 90 sec for 2-3 h, with an exposure time 100 msec for the 488 nm laser. Single z-plane images were acquired with focus on SunTag-GFP foci on the plasma membrane. For experiments in which the GFP fluorescence intensity of individual 24xSunTag arrays was measured, the cells were transfected with a plasmid encoding the 24xSunTag-CAAX protein.

### mRNA sequencing (mRNA-seq) and ribosome profiling (Ribo-seq)

For mRNA sequencing in Fig. 3A and Suppl. Fig. 3A, RNA was isolated from RPE1 *TP53*^KO^ and RPE1 *TP53*^KO^ *EIF2A*^KO^ cells during log-phase growth using an RNeasy Mini kit (Qiagen) according to manufacturer’s instructions. 25 * 106 paired-end reads of 75 bp per sample from a NovaSeq 6000 were aligned to the Hg19 genome build using STAR, to obtain gene-level counts. Genes with low read counts were removed based on the optimal Jaccard similarity index computed using the R package HTSFilter^89^, and differentially expressed genes were identified using edgeR^90^.

For mRNA sequencing in Suppl. Fig. S6A-B, total RNA for mRNA-seq was prepared from RPE1 *TP53*^KO^ *PAC*^KO^ cells harvested after 24 or 48 h of treatment with 250 nM AZD1775 or DMSO in biological duplicate using DirectZol RNA MiniPrep kits (Zymo Research) according to the manufacturer’s instructions. cDNA libraries were generated using TruSeq Stranded Total RNA library kits with Ribo-Zero Gold rRNA depletion (Illumina) and sequenced as 300 nucleotide paired end reads on a NextSeq500 platform (LTRI NBCC). mRNA-seq reads were processed by alignment with Salmon to the reference transcriptome (human genome assembly GRCh38.p13 cDNA)^91^. Differential gene expression changes were calculated with DESeq2 with an FDR cutoff < 0.1 for gene significance^92^. Gene set enrichment analyses were first performed with clusterProfiler^93^. Gene set enrichment analysis (GSEA) software was used to identify the enrichment of genes transcriptionally regulated by AZD1775 (mRNA-seq) in genes identified by ribosome profiling. For Ribo-Seq profiling in Fig. 6A-E and Suppl. Fig. 6D-E, RPE1 *TP53*^KO^ cells were treated with 1 µM of AZD1775 for 24 h. Ribo-seq was performed as detailed in the Supplemental Methods.

## Data availability

The mass spectrometry proteomics data have been deposited to the ProteomeXchange Consortium via the PRIDE partner repository with the dataset identifier PXD054741. The single cell EdU-Seq data have been deposited to the NCBI-GEO repository with identifier GSE287601. Sequencing data of the HAP1 screen is available in the NCBI Sequence Read Archive (SRA) with BioProject number PRJNA1152540. The CRISPR screen data is deposited at the NCBI SRA with BioProject number PRJNA1233308. RNA sequencing data of RPE1 *TP53^KO^* control and *EIF2A*^KO^ cells has been deposited in the Gene Expression Omnibus under accession numbers GSE286143 and GSE291445. Raw sequencing Riboseq data, along with metadata, and count tables have been made available in the Gene Expression Omnibus under the accession number GSE291029.

## Code availability

Scripts for RNA-seq and Ribo-seq library QC evaluation and to produce figures are available on GitHub at https://github.com/mvanins/WEE1_manuscript. The code for analysis and plotting of sc-EdU-Seq data is available on GitHub: https://github.com/jervdberg/WEE1.

## Author contributions

S.d.J, T.R.B, D.D., M.A.T.M.v.V. conceived the project. R.B.T., T.N., M.R., M.L., C.B., M.E., F.J.B. and M.E. performed wet lab experiments. N.H. and T.N. carried out the CRISPR screens that were analyzed by LH. D.B. and T.R.B. performed and analyzed the HAP1 insertional mutagenesis screen. J.Y. prepared purified GCN2 under supervision of F.S. S.Y. and M.E.T. performed and analyzed the socRNA experiments. M.V., A.v.O. and J.v.d.B., performed and analyzed scEdU-seq and RiboSeq experiments. H.R.d.B. analyzed the SILAC proteomics data. R.B.T., T.N., M.R., D.D. and M.A.T.M.v.V wrote the manuscript. All authors provided input on the manuscript.

## Acknowledgements

We thank members of the Durocher and van Vugt lab for critical input. We thank Repare Therapeutics for providing the PKMYT1 inhibitor RP-6306, and Prof. Sir Stephen Jackson for sharing unpublished data and coordination of publication. We thank Henrique Melo for initial CRISPR screen analyses as well as Krzysztof Szkop, Shannon Mclaughlan, Ivan Topisirovic, Ola Larsson for advice and help with polysome profiling and a RiboSeq experiment that was not included in this final manuscript; Kin Chan and Monica Hasegan of the NBCC at the LTRI for expert assistance with sequencing and high content microscopy, respectively. We thank Maarten Hekkelman of the NKI for help with bioinformatics analysis of HAP1 screens. We thank Lisa Koob for assistance with the AZD1775 negative selection CRISPR screen. We thank Claus Sorensen for constructive discussions.

