## Supplemental Information for "WEE1 inhibitors trigger GCN2-mediated activation of the integrated stress response"

This file includes:

Supplementary methods

Supplementary table legends

Supplementary Figure S1

Supplementary Figure S2

Supplementary Figure S3

Supplementary Figure S4

Supplementary Figure S5

Supplementary Figure S6

Supplementary Figure S7

Supplementary Figure S8

#### Supplementary methods

##### Ribosome profiling (Ribo-Seq)

Ribosome profiling libraries were prepared using a version of scRibo-seq[1] that was modified to accommodate higher cell inputs. First, all reaction volumes were scaled up 50× (e.g., from 50 nL to 2.5 µL) to allow manual pipetting. Second, the Micrococcal Nuclease (MNase) concentration was increased to compensate for the increased cell concentration. Specifically, cells were collected by trypsinization (TrypLE, Gibco), counted, washed in PBS0, and approximately 2000 were resuspended in 2.5 µL polysome lysis buffer [20 mM Tris pH 7.5, 15 mM $MgCl_2$, 5 mM $CaCl_2$, 150 mM NaCl, 1% Triton X-100, 2 U/µL RNaseIN Plus (Promega), 0.1 mM Cycloheximide (Sigma Aldrich), 0.1 mM Chloramphenicol (Sigma Aldrich)] and incubated on ice. Next, 2.5 µL of Micrococcal Nuclease (MNase, 62.5 U/µL, New England Biolabs) was added to each sample and incubated at 37 °C for 30 min. All remaining steps of footprint generation, small RNA library construction, and purification were performed as described in scRibo-seq[1]. RNA-seq libraries were prepared using a version of VASA-plate[2] modified to accommodate higher cell inputs. First, all reaction volumes were scaled up 50× (e.g., from 50 nL to 2.5 µL) to allow manual pipetting. Second, the concentration of the CEL-seq2/SORT-seq[3,4] primer was increased to reflect the increased cell concentration. Specifically, cells were collected by trypsinization, counted, washed in PBS0, and approximately 2000 were resuspended in 2.5 µL of 2.5 µM SORT-seq primer, and snap-frozen and stored at -80 °C until processing. Cell lysates were thawed and processed through the steps of lysis and fragmentation, end repair and poly-A tailing, reverse transcription, and second strand synthesis as previously described for VASA-plate[2] with a 50× increase in reaction volume. Ribo-seq libraries were sequenced using v3 chemistry on a NextSeq 2000 (Illumina; NextSeq Control Software version 1.7.0.45751; RTA version 4.12.2) with 72 cycles for read 1, 6 cycles for the i7 index read, and 10 cycles for the i5 index read (sample index). RNA-seq libraries were sequenced using v3 chemistry on a NextSeq 2000 (NextSeq Control Software version 1.7.0.45751; RTA version 4.12.2) with 26 cycles for read 1, 106 cycles for read 2, and 6 cycles for the i7 index read.

For Data analysis of Ribo-seq and RNA-seq data, the reference genome and transcript annotations were obtained from Gencode, using human release 34 (GRCh38.p13). The genome was prepared for alignment by masking all tRNA genes and pseudogenes in the chromosome sequences, and including unique mature tRNA genes as artificial chromosomes. These tRNA genes and psueodgenes were identified using tRNAscan-SE (version 2.0.7) using the eukaryotic and vertebrate mitochondrial models. For metagene analyses, a set of canonical transcripts was defined based on the APPRIS annotations, with the longer isoform being selected in cases where multiple primary APPRIS isoforms existed.

Ribo-seq raw reads were processed, aligned, and quantified as described in scRibo-seq[1]. RNA-seq raw reads were first trimmed using cutadapt (version 3.2) with: -U 6 -U -1 –times 5 -m :15 -A GTTCAGAGTTCTACA -A AAAAAAAAAAAAAAA -A TTTTTTTTTTTTTTT -A CCCCCCCCCCCCCCC -A GGGGGGGGGGGGGGG. Trimmed reads were aligned to the reference genome using STARSolo (version 2.7.10a) with the following parameters: --sjdbOverhang 50 --seedSearchStartLmax 10 --alignIntronMax 1000000 --outFilterType BySJout --alignSJoverhangMin 8 --outFilterScoreMin 0 --outFilterMultimapNmax 1 --chimScoreSeparation 10 --chimScoreMin 20 --chimSegmentMin 15 --outFilterMismatchNmax 5. Aligned reads were deduplicated with UMI-tools (version 1.1.2) using --spliced-is-unique --per-gene --per-contig --per-cell. Count tables report the number of unique reads aligning to any annotated exonic sequence for each gene.

Manipulations, statistics, and plotting were performed in R (version 4.3.3) with dplyr (version 1.1.4), tidyr(1.3.1), data.table (1.15.4), Matrix (1.6-5), ggplot2 (3.5.1), cowplot (version 1.1.3), ggpointdensity (version 0.1.0), and ComplexHeatmap (version 2.18.0).

Differential abundance and translation testing was performed using DEseq2 (version 1.42.1). Library size factors were calculated with scran (version 1.30.2). For the Ribo-seq libraries, these size factors were calculated using the tRNA abundances; for the RNA-seq libraries they were calculated using the protein-coding gene counts. Gene set enrichment analysis was performed using fgsea (version 1.28.0) with default parameters against the Reactome gene sets from msigdbr (version 7.5.1). Scripts for RNA-seq and Ribo-seq library QC evaluation and to produce figures are available at https://github.com/mvanins/WEE1_manuscript.

**Single-cell EdU-Seq**

After ligation of fragments to T7 promoter-containing adapters with cell barcodes and UMI, the contents of each plate were collected into VBLOK200 reservoirs precoated with mineral oil by centrifugation. The aqueous phase was collected and separated from any residual mineral oil by centrifugation. EdU–PEG3–biotin containing DNA molecules was affinity purified using MyOne Streptavidin C1 magnetic beads (Invitrogen) according to the manufacturer’s protocol. Subsequently, we retrieved the complementary strand of the EdU–PEG3–biotin containing the DNA strand by heat denaturation at 95°C. While ramping down the temperature (0.1 °C $s^{-1}$) to 20°C, we annealed an oligo (Suppl. Table 3) complimentary to the constant adapter sequence region in oligo annealing buffer (20 mM Tris pH 8, 1 mM $MgCl_2$, 100 mM NaCl). Next, the primer was extended to generate double-strand DNA using Klenow large fragment mix for 45 min at 25°C, and heat-inactivation for 20 min at 75°C. DNA fragments were purified at a sample-to-bead ratio 1:1 and resuspended.

Pre-amplified libraries were linearly amplified using a MEGAscript T7 Transcription Kit (ThermoFisher Scientific) for at least 12 h at 37°C. Template DNA was removed, and amplified RNA (aRNA) was fragmented for 2 min at 94 °C with fragmentation. aRNA was directly cooled to 4 °C and fragmentation was stopped. The fragmented RNA was purified and eluted in 12 µl of H2O. Next, 5 µl of aRNA was converted to cDNA by RT in two steps. First, the RNA was primed for RT by adding 0.5 µl of dNTPs (10 mM) and 1 µl of random hexamer RT primer 20 µM (Suppl. Table 3) at 65°C for 5 min followed by direct cooling on ice. Second, RT was performed by the addition of 2 µl of first-strand buffer, 1 µl of 0.1 M dithiothreitol, 0.5 µl of RNAseOUT and 0.5 µl of Superscript II, and incubating the mixture at 25°C for 10 min, followed by 60 min at 42°C and 20 min at 70°C. Single-strand cDNA was purified from aRNA through incubation with 0.5 µl of RNAseA (ThermoFisher Scientific) for 30 min at 37 °C. Finally, cDNA was amplified by PCR. DNA fragments were purified twice with Ampure XP beads (Beckman Coulter) at a sample to beads ratio 0.8:1, and were resuspended. The abundance and quality of the final library were assessed by Qubit and Bioanalyzer.

**Quantitative image-based cytometry (QIBC)**

For immunofluorescent staining of ATF4, cells were grown on round glass coverslips, no. 1.5 glass bottom 35mm dishes (MatTek), or clear bottom opaque 96-well plates (Corning #3603) for quantitative image-based cytometry (QIBC) and fixed with 4% paraformaldehyde (PFA) in PBS at RT or ice-cold methanol at -20°C for 20 min. For PFA fixation, cells were permeabilized with PBS-0.3% Triton X-100 for 20 min at RT. Blocking was performed for 30 min in 0.1% PBST with 3% BSA or goat serum blocking buffer (10% goat serum, 0.5% NP-40, 0.5% saponin in PBS). Cells were stained with a primary antibody against ATF4 (1:200; Cell Signaling, #11815) in blocking buffer for 2 h at RT. Cells were washed three times with 0.1% PBST and then stained with Alexa Fluor 647 goat anti-rabbit IgG (1:1000; Thermo Fisher, A21244) and 0.4 µg/ml DAPI for 1 h at RT in the dark. For S-phase labeling with 5'-ethynyl-2'-deoxyuridine (EdU, Thermo Fisher) click chemistry, EdU was added to culture media at a final concentration of 20 µM 30 min before harvest. Cells were fixed with PFA a second time after secondary antibody staining and incubated in Click-iT reaction buffer (100 mM Tris HCl pH 8.5, 1 mM CuSO4, 100 mM ascorbic acid, 10 µM Alexa Fluor 488 or 647 azide) for 30 min in the dark. Subsequently, cells were imaged or mounted on glass slides using ProLong Gold antifade reagent (Invitrogen). Slides were imaged on a Zeiss LSM780 confocal microscope with a 63x oil-immersion objective. Cells stained in 96-well plates were imaged on a GE InCell Analyzer 6000 using a 20x air objective. Images were analyzed using Acapella based scripts on the Columbus high-content image analysis platform (PerkinElmer). For QIBC, analysis was first performed using DAPI fluorescence to segment nuclei and then measuring the median or sum fluorescence intensities of each channel (DAPI, EdU or ATF4). To measure chromatin bound RPA2 in mitotic cells, H3S10 fluorescence was used to segment nuclei. To measure fluorescence intensities specifically in S-phase cells, EdU staining intensity histograms were drawn and high EdU incorporation was manually gated. G1 and G2-phase cells were defined by low EdU incorporation and 2n or 4n DNA content (sum DAPI fluorescence), respectively.

**Supplementary Table 1.** mRNA sequencing GSEA Reactome analysis results.

**Supplementary Table 2.** List of synergy scores of the AZD1775, Debio 0123 and RP-6306 synergy assays.

**Supplementary Table 3.** Overview of sgRNA and primer sequences.

**Supplementary Table 4.** List of DrugZ scores.

**Supplementary Table 5.** List of MAGecK scores.

#### Supplemental Figure S1

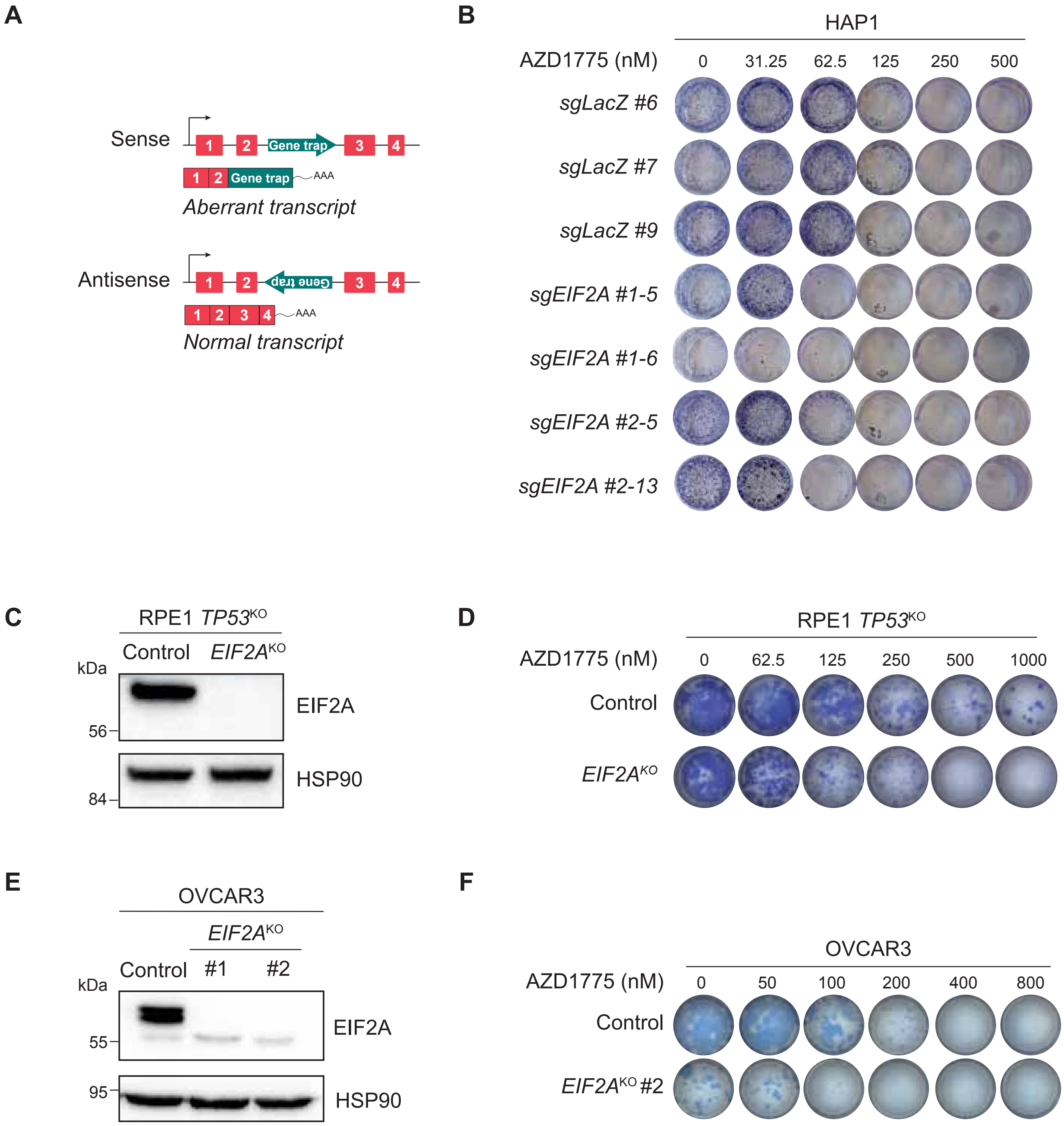


**Supplementary Figure S1. Loss of EIF2A sensitizes cells to AZD1775.** (**A**) Schematic representation of the HAP1 viral insertional mutagenesis screen. Insertion of a gene-trap with a strong splice acceptor in sense orientation results in gene disruption, whereas anti-sense insertion will not. (**B**) Clonogenic survival analysis of HAP1-sgLacZ cells and HAP1-sgEIF2A clones treated with AZD1775. (**C**) Immunoblot analysis of RPE1 *TP53*$^{KO}$ control and *EIF2A*$^{KO}$ cells. (**D**) Representative images of RPE1 *TP53*$^{KO}$ and RPE1 *TP53*$^{KO}$ *EIF2A*$^{KO}$ cells treated with AZD1775 at indicated doses for 10 days. (**E**) Immunoblot analysis of OVCAR3 control and *EIF2A*$^{KO}$ #1 and #2 cells. (**F**) Representative images of OVCAR3 control and OVCAR3 *EIF2A*$^{KO}$ #2 cells treated with AZD1775 at indicated doses for 10 days (n=1 biological replicate).

#### Supplemental Figure S2

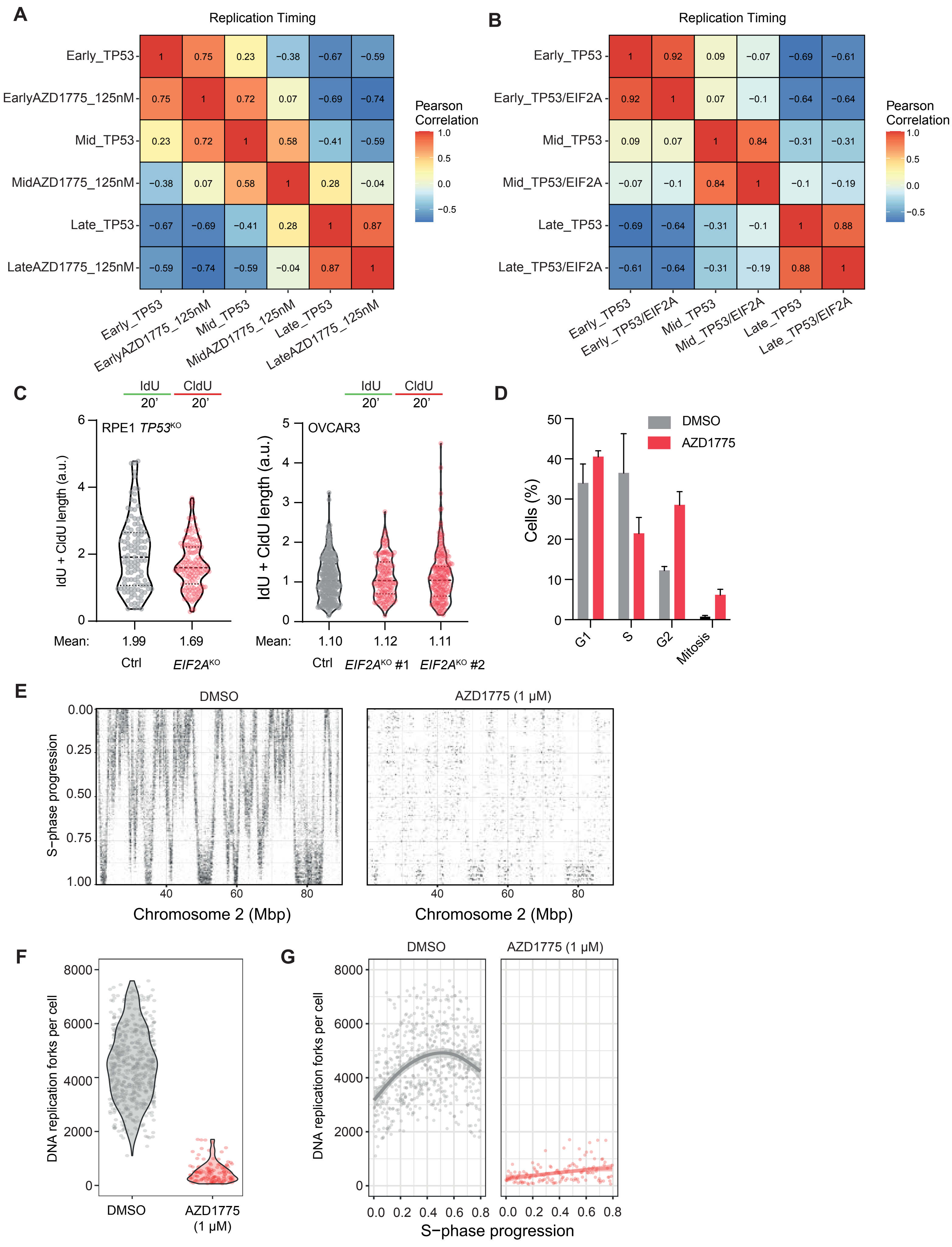


**Figure legend on following page**

#### Supplemental Figure S3

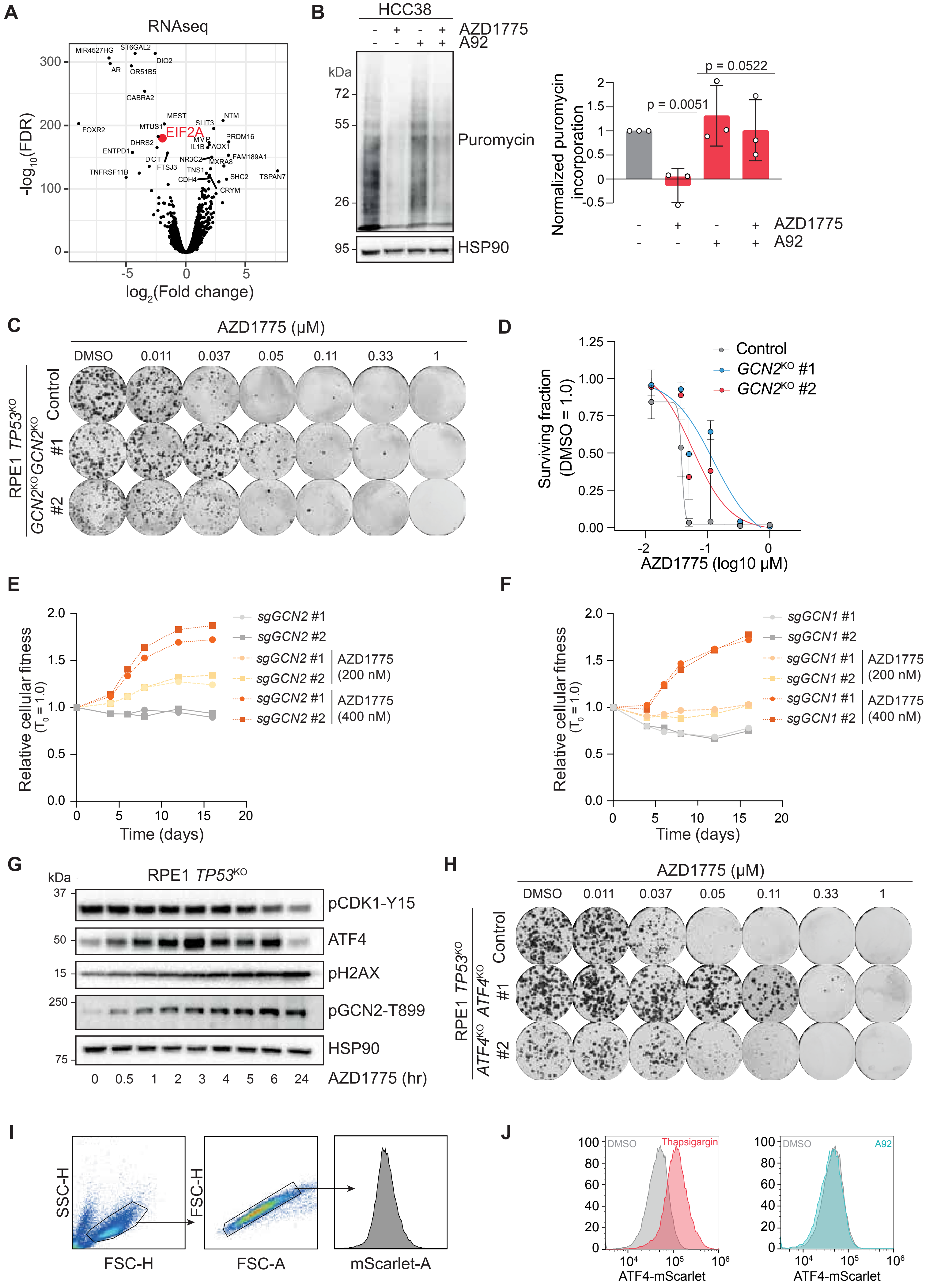


**Figure legend on following page**

#### Supplemental Figure S4

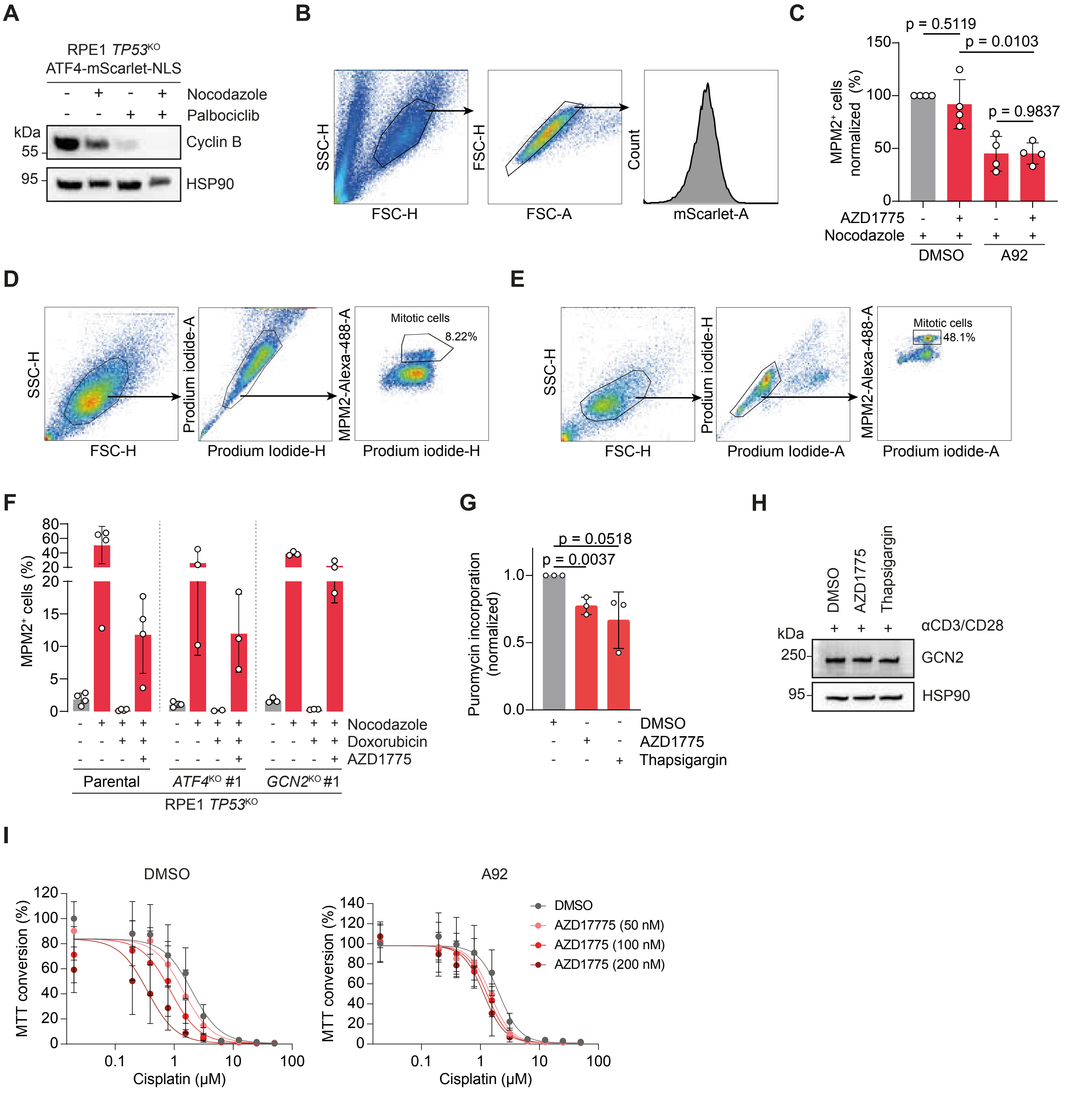


**Supplementary Figure S4. Integrated stress response activation by WEE1i is cell cycle independent.** (**A**) Immunoblot analysis of MPM2 in RPE1 *TP53*KO ATF4-mScarlet-NLS cells after treatment with nocodazole and doxorubicin as indicated in Fig. 4B. (**B**) Gating strategy for flow cytometry analysis shown in Fig. 4C. (**C**) Flow cytometry analysis of MPM2 in RPE1 *TP53*KO cells after treatment with AZD1775 (1 µM) and/or A92 (1 µM) in the presence of nocodazole. Data represent mean ± SD (n=4). Statistical analysis was performed using an unpaired t-test with p≤0.05 considered significant. (**D**) Gating strategy for flow cytometry analysis in Fig. 4F. (**E**) Gating strategy for flow cytometry analysis in Fig. 4G and Suppl. Fig. 4F. (**F**) Flow cytometry analysis of MPM2-positivity in RPE1 *TP53*KO *PAC*KO control or *ATF4*KO #1 or *GCN2*KO#1 cells after treatment with nocodazole, doxorubicin (0.5 µM) and AZD1775 (1 µM). Mean ± SD (n=4 for all parental conditions, n=3 for all KO conditions except for *ATF4*KO #7 nocodazole + doxorubicin, n=2). (**G**) Quantification of puromycin incorporation assay shown in Fig. 4H. Mean ± SD (n=3). Statistical analysis was performed using an unpaired t-test with p≤0.05 considered significant. (**H**) Immunoblot analysis of PBMCs stimulated with anti-CD3/CD28 beads. PBMCs were treated with DMSO, AZD1775 (500 nM) or thapsigargin (500 nM) for 24 h. (**I**) MTT conversion of RPE1 *TP53*KO cells after treatment with the indicated doses of cisplatin and AZD1775 or A92 (200 nM) for 5 days.

#### Supplemental Figure S5

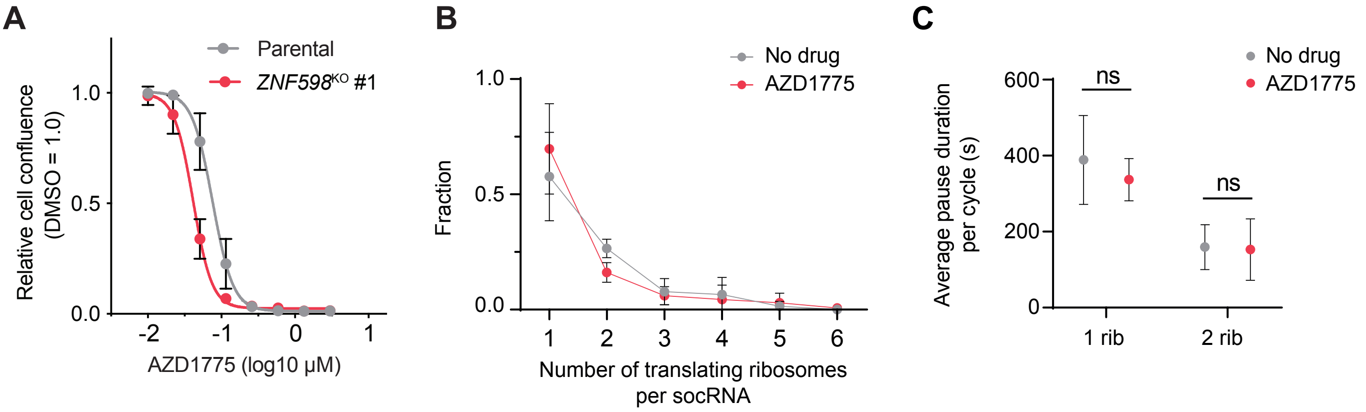


**Supplementary Figure S5. Effects of WEE1i on translation kinetics.** (**A**) Cell confluence analysis of parental RPE1 *TP53*$^{KO}$ *PAC*$^{KO}$ or *ZNF598*$^{KO}$ #1 cells after treatment with indicated doses of AZD1775. Mean ± SD, n=3 technical replicates. (**B**) Number of translating ribosomes per socRNA in U2OS cells after treatment with AZD1775 (300 nM, 24 h). Data represent mean ± SD (control n=2; AZD1775 n=3). (**C**) Average pause duration on socRNA with Xbp1(S255A) pause sequence in U2OS cells after treatment with AZD1775 (300 nM, 24 h). Data represent mean ± SD (No drug n=2; AZD1775 n=3).
reporter cells afer treatment with thapsigargin (1 μM) or A92 (1 μM) for 24 h, measured by flow cytometry.

#### Supplemental Figure S6

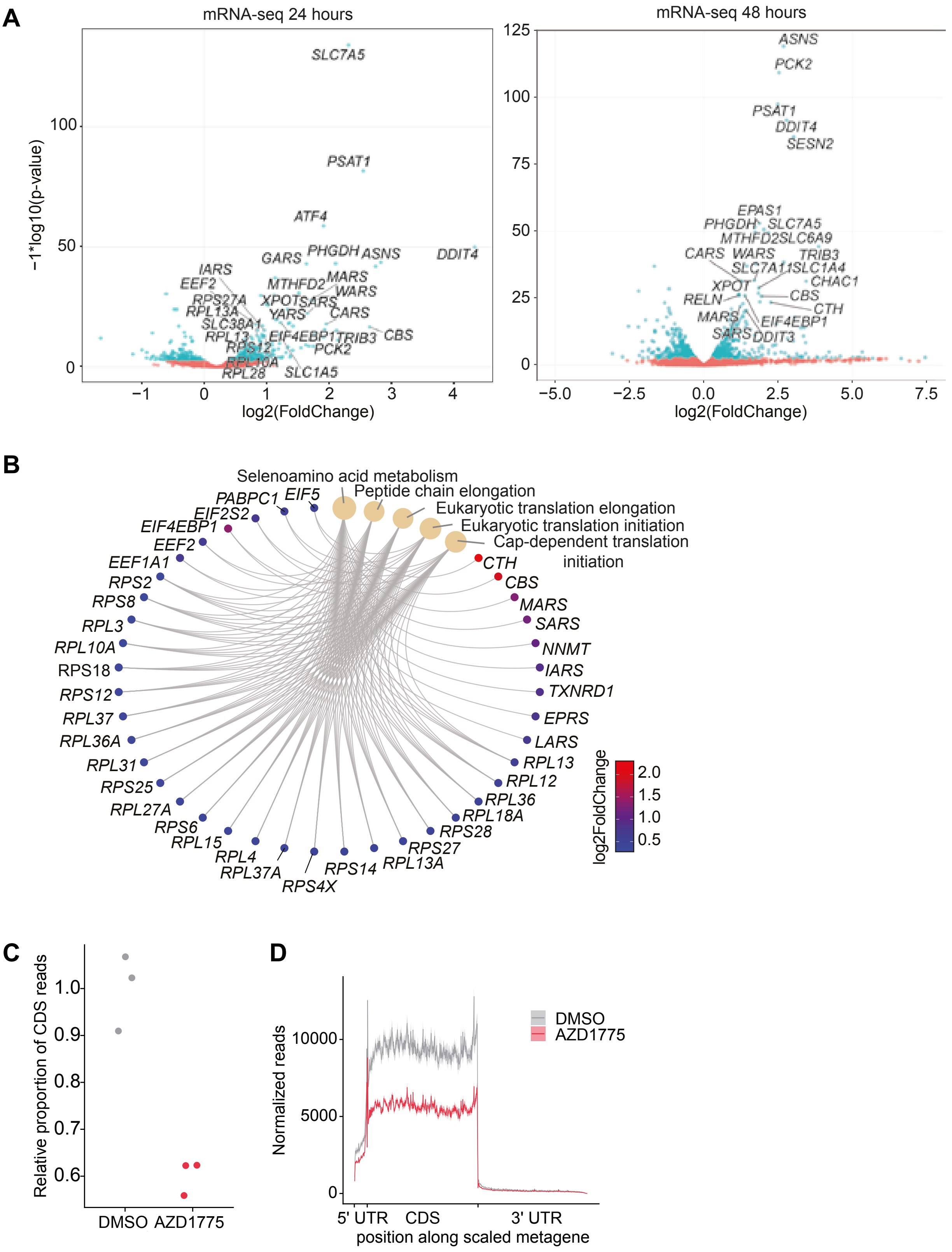


**Supplementary Figure S6. Differential gene expression analysis upon AZD1775 reveals increased expression of translation initiation and elongation genes.** (**A**) Volcano plot showing significantly DEGs (blue points) after 24 h (left) or 48 h (right) of treatment with AZD1775 (250 nM) relative to DMSO. (**B**) GSEA performed with differentially expressed genes (DEGs) from RPE1 *TP53*$^{KO}$ *PAC*$^{KO}$ cells shown in Fig. 5D (AZD1775, 48 h). (**C**) Relative proportion of coding DNA sequence reads after treatment with vehicle or AZD1775 in RPE1 *TP53*$^{KO}$ cells (n=3). (**D**) Normalized reads after treatment with vehicle or AZD1775 in RPE1 *TP53*$^{KO}$ cells.

### Supplemental Figure S7

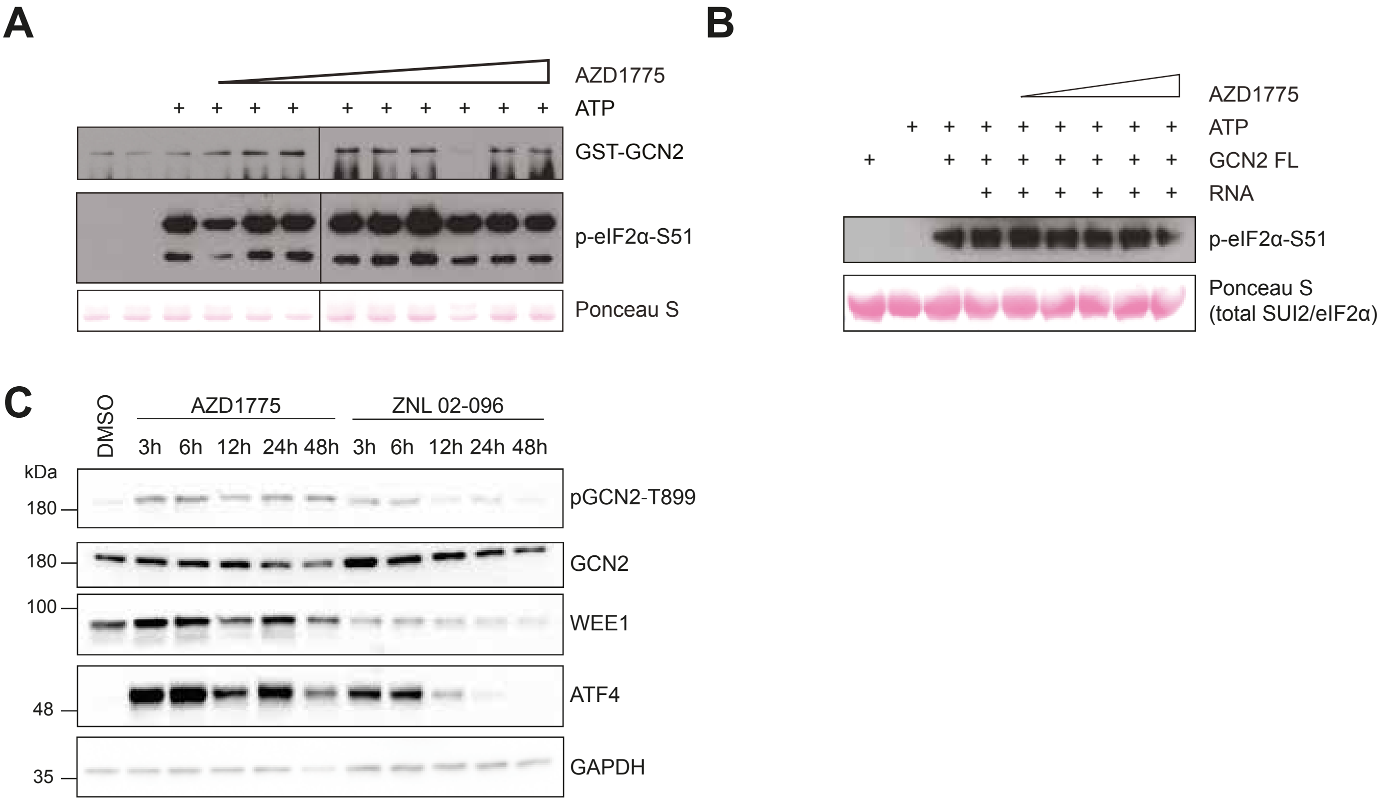


**Supplementary Figure S7. The WEE1i AZD1775 does not directly interact with the GCN2 kinase domain.** (**A**) Kinase assays with recombinant yeast eIF2α (SUI2) and GST-tagged purified GCN2 kinase/pseudokinase domain with 1-100 nM concentrations of AZD1775. (**B**) Kinase assays performed with recombinant SUI2, total RNA from HEK293T cells, and human full-length GCN2 purified from HEK293T. AZD1775 was titrated into reactions to final concentrations of 1-500 nM. (**C**) Immunoblot of RPE1 *TP53*$^{KO}$ cells treated with AZD1775 (100 nM) or ZNL 02-096 (250 nM) for the indicated times.

#### Supplemental Figure S8

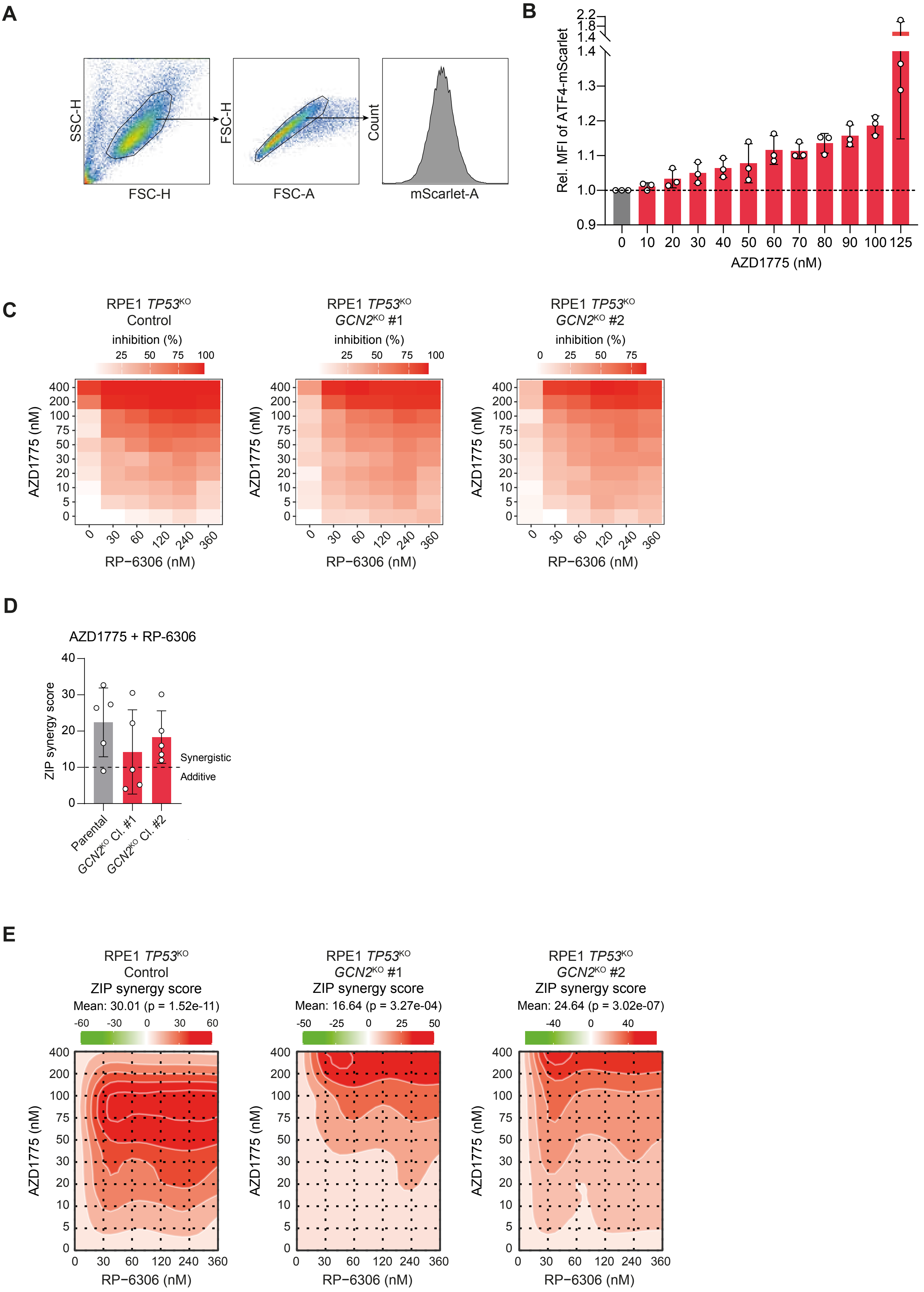

**Supplementary Figure 8. PKMYT1i and WEE1i synergistically induce cytotoxicity without integrated stress response induction.** (**A**) Gating strategy for flow cytometry analyses shown in Fig. 8A, E, and Suppl. Fig. 8B. (**B**) Relative ATF4-mScarlet MFI in RPE1 ATF4-mScarlet reporter cells after treatment with the indicated doses of AZD1775. Data represent mean ± SD (n=3). (**C**) Cell viability analysis in parental RPE1 *TP53*$^{KO}$ *PAC*$^{KO}$, *GCN2*$^{KO}$ #1 and *GCN2*$^{KO}$ #2 cells after treatment with AZD1775 and RP-6306 at the indicated doses (n=5). (**D**) ZIP synergy scores of cell survival matrices shown in Suppl. Fig. 8C and 8E. Data represent mean ± SD (n=5). (**E**) Synergy plots with ZIP synergy scores of heatmaps shown in Suppl. Fig. 8D.
